# ELIMÄKI locus is required for mechanosensing and proprioception in birch trees

**DOI:** 10.1101/616474

**Authors:** Juan Alonso-Serra, Xueping Shi, Alexis Peaucelle, Pasi Rastas, Matthieu Bourdon, Juha Immanen, Junko Takahashi, Hanna Koivula, Gugan Eswaran, Sampo Muranen, Hanna Help-Rinta-Rahko, Olli-Pekka Smolander, Chang Su, Omid Safronov, Lorenz Gerber, Jarkko Salojärvi, Risto Hagqvist, Ari-Pekka Mähonen, Kaisa Nieminen, Ykä Helariutta

## Abstract

The remarkable vertical and radial growth observed in tree species, encompasses a major physical challenge for wood forming tissues. To compensate with increasing size and weight, cambium-derived radial growth increases the stem width, thereby supporting the aerial body of trees. This feedback appears to be part of a so-called “proprioception” (^1, 2^) mechanism that controls plant size and biomass allocation. Yet, how trees experience or respond to mechanical stress derived from their own vertical loading, remains unknown. Here, we combined two strategies to dissect the proprioceptive response in birch. First, we show that in response to physical loading, trees promote radial growth with different magnitudes along the stem. Next, we identified a mutant cultivar (*B. pubescens cv. Elimäki*) in which the main stem shows normal vertical development, but collapses after three months. By inducing precocious flowering, we generated a backcrossed population (BC_1_) by producing two generations in 4 years. In his scheme, we uncovered a recessive trait (*eki*) that segregates and genetically maps with a Mendelian monogenic pattern. Unlike WT, *eki* is resistant to vertical mechanical stimulation. However, *eki* responds normally to the gravitropic stimulus by making tension wood. Before the collapse, cell size in *eki* is compromised resulting in radial growth defects, depending on stem height. Cell walls of developing xylem and phloem tissues have delayed differentiation in *eki*, and its tissues are softer compared to WT as indicated by atomic force microscopy (AFM). The transcriptomic profile of *eki* highlighted the overlap with that of the *Arabidopsis* response to touch. Taken together, our results suggest that the mechanical environment and cell wall properties of developing woody tissues, can significantly affect the growth responses to vertical loading thereby compromising their proprioceptive capacity. Additionally, we introduce a fast forward genetics strategy to dissect complex phenotypes in trees.

## Introduction

Plant development involves a continuous change in size and shape. Since the emergence of land plants (≈ 470 mya) a terrestrial environment introduced physical challenges that combined with individual genotypes shaped a new diversity of plant growth habits. During this process, various plant architectures succeeded in the competition for light and pollinators while resisting strong winds, rain and snow. In this junction of evolution and development, highly specialized vascular cells and tissues were positively selected to perform functions of support and transport. Due to their large size, trees are particularly specialized in these characteristics making them an interesting case study. In tree species, while lifting up the aerial plant body is crucial for plant life, growth proportions or plant allometric features (e.g stem height vs. diameter), must be maintained below the threshold that compromise the physical stability (3). Therefore, sensing these mechanical constraints through local tissue deformations, and consequently modulating growth to adjust the stem stability, represents an important developmental feedback to provide plants with postural perception. This is part of mechanism known as “proprioception” (1, 4). The apical growth of trees is controlled by the cell division rate in the shoot apical meristem (SAM) that produces axillary primordia (later named stem nodes), combined with the longitudinal extension of shoot internodes. On the other hand, radial growth is achieved by the activity of vascular cambium (VC), a cylindrically shaped meristem that increases the stem width by producing xylem tissues inwards and phloem tissues outwards. Given that most (woody) tissues in an adult tree are produced by the VC, this meristem has a central role in providing the final long-term physical properties of the stem. Interestingly the mechanics associated with cambial activity and radial growth remain largely understudied, partly due to the inaccessibility of this meristem. How do compressive forces derived from increasing plant body weight integrate with the internal expansive forces derived from the meristem activity? Ultimately, do these forces modulate how trees grow?

A positive correlation between aerial biomass (kg) and stem diameter has been measured in multiple species (5), and growth is partially controlled by mechanical stress in self-supported plants such as trees (6–8). Mechanical treatments like stem bending or tilting induces gravitropism, stem asymmetric thickening, and later reaction wood formation (9, 10). In these conditions a proprioceptive mechanism seems to control the straightening of stems (2), but here the stem experiences different orientations (respect to the gravity vector) during the treatments, therefore differing mechanically from the solely vertical loading. In the vertically standing plants, external support produces taller and thinner stems (11) and experiments with *Arabidopsis thaliana* revealed that inflorescence stems can react to externally added weight by increasing xylem differentiation (12). Moreover, the mechanical pressure in the VC may be required for xylem differentiation in trees (13). This suggest that the VC responds to mechanical stimulus by integrating multi-scale signals derived from vertical loading and radial growth (maturation stress) (14). Interestingly, the direct response of tree stems to a vertical static loading on was not tested before. At the cellular level, apical meristems in *Arabidopsis* respond to different mechanical stress patterns, which in turn contribute to the control of meristem activity (15–17). In addition, mechanical properties of cell walls are sufficient to promote or restrict the developmental progression (16). Also, the study of cell wall mutants has exposed the pre-existing physical forces in plant tissues (18). These examples highlight the power of genetic manipulations of the mechanical environment to learn from the subsequent developmental outputs. Additionally, dissecting the implications of global and local mechanics requires a high-resolution approach to organs, tissues and cells.

While in *Arabidopsis* is possible to easily identify highly informative mutants, the same approach is hindered in trees by the lack of mutant collections, or their long generation time. Stem mechanics and architecture have been key traits of interest in plant breeding since the Green Revolution, but, due to their long generation time, forward genetics approach in forest tree species remains underdeveloped. Birch (*Betula spp.*), an emerging model tree, is among the few reported tree species where flowering can be reduced from 7-10 to 1 year under specific greenhouse conditions (19). In addition, *Betula* species are diverse in their stem architecture exhibiting bushy forms, dome-shaped crowns and normally straight standing stems. By creating a collection of these cultivars we have previously identified a stem architecture mutant of *B. pendula* carrying a homozygous mutation in LAZY1, which in *Arabidopsis* is responsible for gravitropism (20, 21). Therefore, this features make birches an ideal model to screen for developmentally affected genotypes and after segregating the traits of interest, study the developmental implications in loss-of-function scenarios.

Here, we first tested the proprioceptive response of WT trees to vertical manipulations of stem weight, and revealed that trees respond by promoting growth in different magnitudes. Next, we studied a naturally occurring mutant of downy birch (*Betula pubescens*) called *Elimäki* (*eki*). The *eki* trees have a distinctive trait, in which the main stem mechanically fails to support trees and collapses after 3 months of normal vertical growth. By implementing induced precocious flowering we produced a BC_1_ segregating population in 4 years. The genetic analysis and QTL mapping of the segregating *eki* phenotype fits a Mendelian pattern consistent with a single recessive locus. Before collapse, *eki* trees are resistant to the same treatments used before in WT indicating that the EKI locus is required for proprioceptive responses. Then, an integratory analysis of tissues transcriptome, morphology, composition and mechanical properties uncovered the developmental components of the mutant phenotype. We conclude that the mechanical environment of wood forming tissues can significantly impact on the proprioceptive responses of trees. At a transcriptional level, the overlapping stress pattern between *eki* and the *Arabidopsis* touch response suggest the involvement of mechanosensing as a major contributor to proprioception in birch.

## Results

### Birch trees respond to vertical manipulations of stem weight

In addition to stem height and diameter being affected by natural or artificial bending, the stem thickness of standing trees also positively correlates with their crown biomass (5). Thus, since secondary growth is an additive process, we tested if vertical manipulations impacting the stem weight, are sufficient to affect the stem diameter and/or height. Since there were no published examples of this experiment, we designed four different scenarios and compared the results between them (**Fig. 1A**): Scenario 1: non-treated supported trees (no extra weight added), Scenario 2: added weight on supported trees, Scenario 3: added weight on partially supported trees (permitting small stem movements), and Scenario 4: vertically pulled of stems from the top. The reasoning for scenario 3 was to test whether stem movements would be required for proprioceptive responses. While in scenarios 2 and 3 the weight effect was directed downwards, in scenario 4 the effect was upwards. The treatments were constant and applied gradually along the upper half of the stem to imitate a natural weight distribution. Finally, we used the trees in scenario 1 as a control of growth without vertical weight manipulations. To provide a holistic interpretation of the changes in stem proportions we measured: stem height, internode number, and diameters (D). Diameters were measured in 7 vertical positions: D1 to D4 (last 4 internodes in the base), D5 (50% height internode), D6 (62.5% height internode) and D7 (75% height internode) (**Fig. 1E**). Finally, all parameters were documented by 4 time points (T1-T4) in 3 weeks.

**Figure 1.**
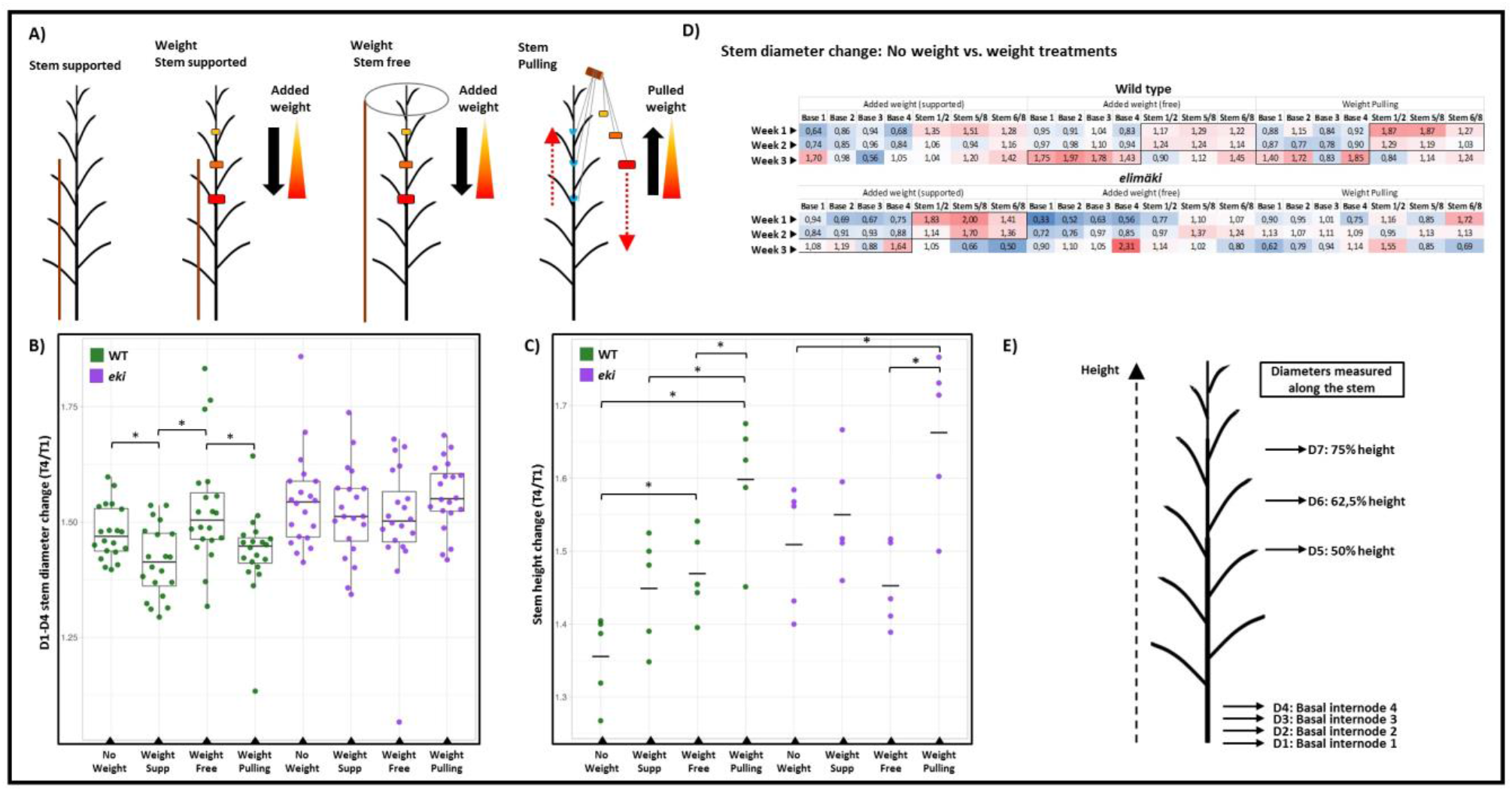
Vertical mechanical manipulations in WT vs. *eki*. **A)** Experimental setup for vertical mechanical manipulations of stem weight. Images show: Scenario 1 (no weight added), Scenario 2 (weight added on supported stems), Scenario 3 (weight added on semi-free stems), Scenario 4 (stems pulled up from the top), red arrows indicate the pulling direction. **B)** Stem diameter change in final vs. initial time-points (T4/T1) on the last 4 internodes from the base (D1-D4). **C)** Stem height change (T4/T1) from the same trees shown in B. **D)** Response ratio in weight vs. no weight treatments (Scenario 1), ratios > 1 shows higher AVG response in Scenario 2, 3 or 4 compared to 1. The table also indicates the increments per week (e.g. week 1 = T2/T1) and the different stem position where diameters were measured. **E)** Diagram of the measured stem position. Each experimental setup consisted of 5 trees per treatment. Asterisks indicate statistically significant differences (*p*≤0.05) by Student-t test.

Taken together the various treatments, we observed that manipulating trees with vertical stress results in enhanced stem diameter. We found that the proportional increments of stem diameter (final/initial or T4/T1) were promoted in the scenario 3 as compared to the scenarios 1, 2 and 4. In the basal internodes (D1-D4), the differences were significantly larger in scenario 3 vs. 2 and 4 (**Fig. 1B**). When combining the middle (D1-D5) or all measured internodes (D1-D7), scenario 3 was also larger than scenario 1 (**Fig S1**). These results suggest that freedom of stem movement may be required for weight-induced responses of secondary growth.

The analysis of diameter changes along the stem (D1-D7), highlights that within the same time period, upper sections (D5-D7) underwent a larger proportional change of thickness as compared to the basal internodes (D1-D4) (**Table S1**). Interestingly, when the scenarios 2, 3 and 4 are compared against scenario 1 (Ratio: weight treatments/no weight), the time series revealed that the first response occurred in the upper sections of the stem and a later one at the base. This pattern was clearer in Scenarios 3 and 4 (**Fig 1D**). Finally, we found an increase of height across treatments where the pulling scenario had the strongest effect (**Fig. 1C**). The changes of internode numbers were similar across all scenarios (**Fig S1**). Combined these results indicate that free-standing WT trees respond to the stimuli of vertically applied weight. This response occurs by adjusting the pattern of radial growth in different magnitudes along the stem. In addition, while height changes were different between scenarios, the internode numbers remained similar; indicating that this mechanism may be independent from the meristematic activity in the shoot apex, but linked to the activity of the vascular cambium.

### Forward genetics in birch identifies a recessive trait linked to stem mechanics

The response of tree stems to mechanical stimulus is a complex multifactorial phenomenon. To genetically dissect this trait, we created a collection of birch cultivars with defective proprioceptive or architectural phenotypes and performed inheritance analysis by selfing and outcrossing these individuals (**Fig 2A**). In this first step, we selected segregating traits with recessive genetic inheritance (year 1). Among others, we identified a mutant named after its original location as “*Elimäki Original*” (EO). EO has an abnormal stem growth pattern, the original tree has a high number of irregular branches. By combining genome sequencing and flow cytometry based ploidy analysis, we confirmed that EO is in the tetraploid background and genetically close to *B. pubescens* (4n = 56) (**Fig S2-S3a and Salojärvi et al., 2017**). The main stem of self-pollinated EO progenies (S_1_), collapsed within three months of growth (**Fig. 2A**); instead, seedlings from wind-pollinated flowers (F_1_) produced normal, upright standing trees, confirming that the *Elimäki* trait (*eki*) is recessive. After inducing early flowering in a 2 years-old F_1_ individual, we backcrossed it with EO to generate a BC_1_ population. In this population of 60 individuals the segregation of standing vs. collapsing trees fitted the 1:1 segregating scenario (**Table S2**). Trees in BC_1_ with the *eki* trait were not able to support their stem longer than 3 months. The ploidy analysis of 10 individuals BC_1_ (5 WT and 5 *eki*) confirmed their tetraploidy (**Fig S2**). These results showed that the *eki* phenotype is genetically inherited and segregates in the pattern of a Mendelian monogenic locus in the tetraploid background.

**Figure 2:**
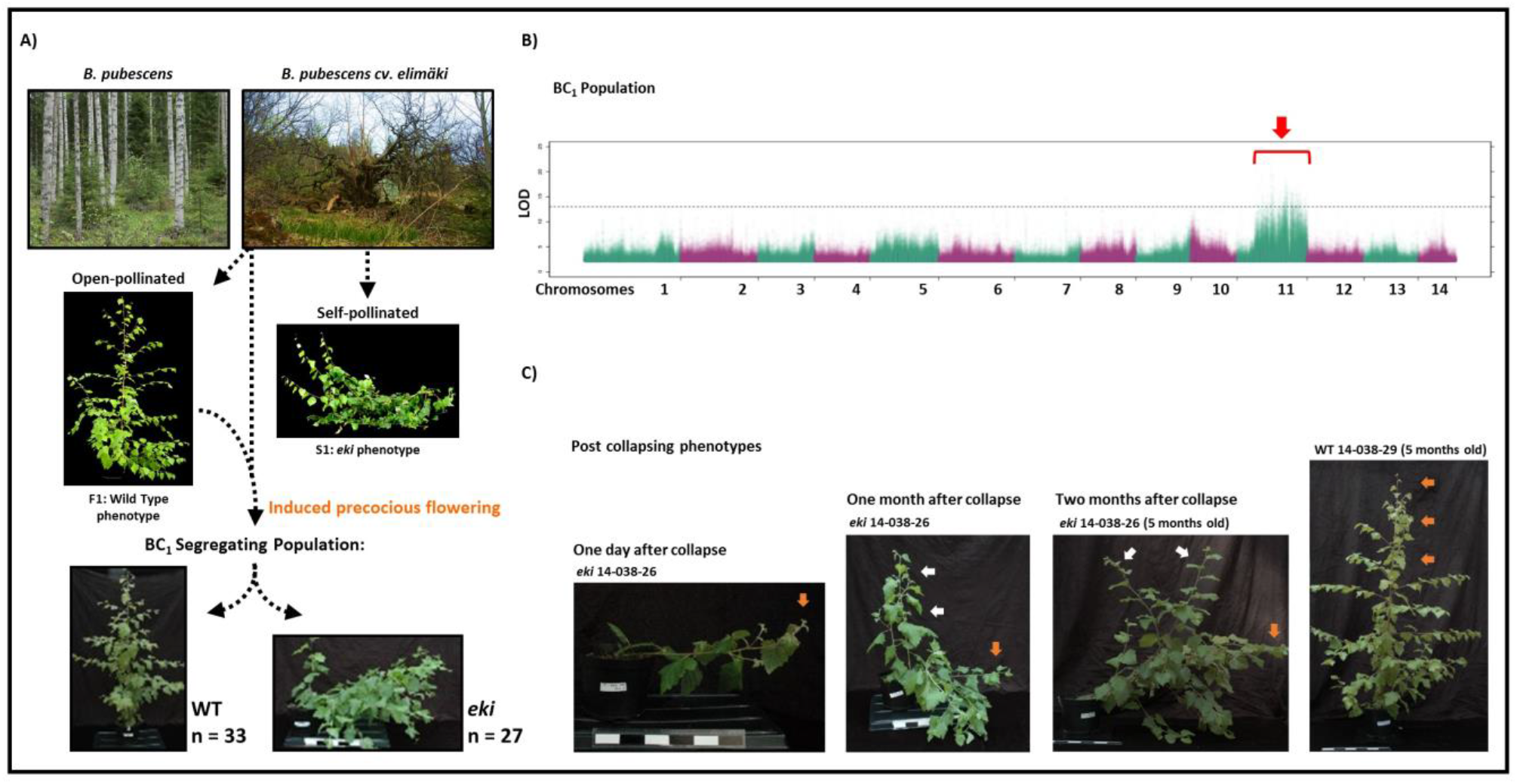
Forward genetics approach identified a stem mechanics mutant. **A)** Crossing scheme to obtain the BC_1_ segregating population: the naturally occurring mutant *B. pubescens cv. Elimäki (EO)*, has a distinctive stem architecture with multiple dominant and irregular branches. The progeny of self-pollinated trees (S1), resembled that of EO, whereas open-pollinated F1 individuals rescued the WT phenotype. F1 was induced to flower and backcrossed with EO to produce BC_1_ **B)** Manhattan Plot showing the association of polymorphic markers to WT vs. *eki* phenotypes, identified in the BC_1_ population by separately pooling WT and *eki* data. Candidate mapping window is indicated by a red arrow **C)** Phenotype progression of the *eki* phenotype. *eki* trees showed gravitropic response one day after collapse, on the following month the outgrown branches successively collapsed *Betula pubescens* picture was provided by Erkki Oksanen (Luke-Finland).

In order to identify a genetic association with the *elimäki* phenotype we sequenced individually 60 trees from the BC_1_ population and its parents EO and F1 with higher depth. This sequencing data was mapped to *B. pendula* genome (20) and a linkage map with about 2.5 million simplex (AAAB x AAAA) markers and 56 linkage groups (likely 4n=56) were constructed using Lep-MAP3=LM3 (22) and new pipeline to utilize polyploid species. Each *B. pendula* chromosome corresponded to four linkage groups of 56 in *B. pubescens* (**Fig S3b**). The QTL mapping was conducted by calculating LOD score between segregation pattern of markers (imputed by LM3) and the binary *eki*/WT phenotype. The end of chromosome 11 gave the most significant association both in the BC_1_ population (**Fig 2B**) and to one of EO parent’s linkage group with a LOD score of over 5 (hamming distance of 12/60, p<0.009 (two-sided binomial distribution + bonferroni correction with the number of different markers = 2749) (**Fig S4**). The identified mapping window in chromosome 11, contains 324 annotated genes in *B. pendula* genome (**Supplementary Data 1**). Further analysis using RNASeq from stem tissues identified that 76 of these genes are differentially expressed in *eki* compared WT, and the expression of 23/76 is affected in all stem samples (**Supplementary Note 1**).

### The *eki* trees respond to gravity, but do not respond to vertical mechanical stimulations

The phenotype characterization in the BC_1_ population showed that the collapsing phenotype occurred within three months after germination in a narrow time window (**Fig 2B, Supplementary Data 2**). This took place through a distinct bending at the base of the main stem. In most cases the trees fell down rapidly, within 24 hrs. One day after the collapse the shoot tips and branches showed gravitropic response, on the following month initiation of side-branches was triggered near the base of the stem. While these branches initially displayed a vertical orientation, their ability to stand upright was eventually compromised resulting in successive collapse, and finally leading to a highly branched bushy-like architecture (**Fig 2C**). After the collapse, the main stem was able to produce tension wood in response to the gravity stimulus, indicating that the gravitropic perception and response were not affected by the *eki* mutation (**Fig S5**). Before collapse the stem height was similar between the WT and *eki* trees (**Fig. 3B**), and *eki* had in average 2,5 internodes more than WT (**Fig S5C**) indicating that primary growth is not compromised in *eki*. In contrast, in 60 days old trees radial growth was significantly reduced towards the base of the stem in *eki* just before the collapse (**Fig. 3C-E**). However, when the stem diameter progression was analyzed in time and at fixed position (10cm height), *eki* was significantly thinner than WT only after 60 days of growth, and not before (**Fig S5B**). Therefore, these results indicate that secondary growth in *eki* was progressively compromised as the tree size increased. To study this, we explored whether the diameter reduction in *eki* could be explained through the lack of adjusted growth response to the mechanical stress caused by increasing tree weight.

**Figure 3.**
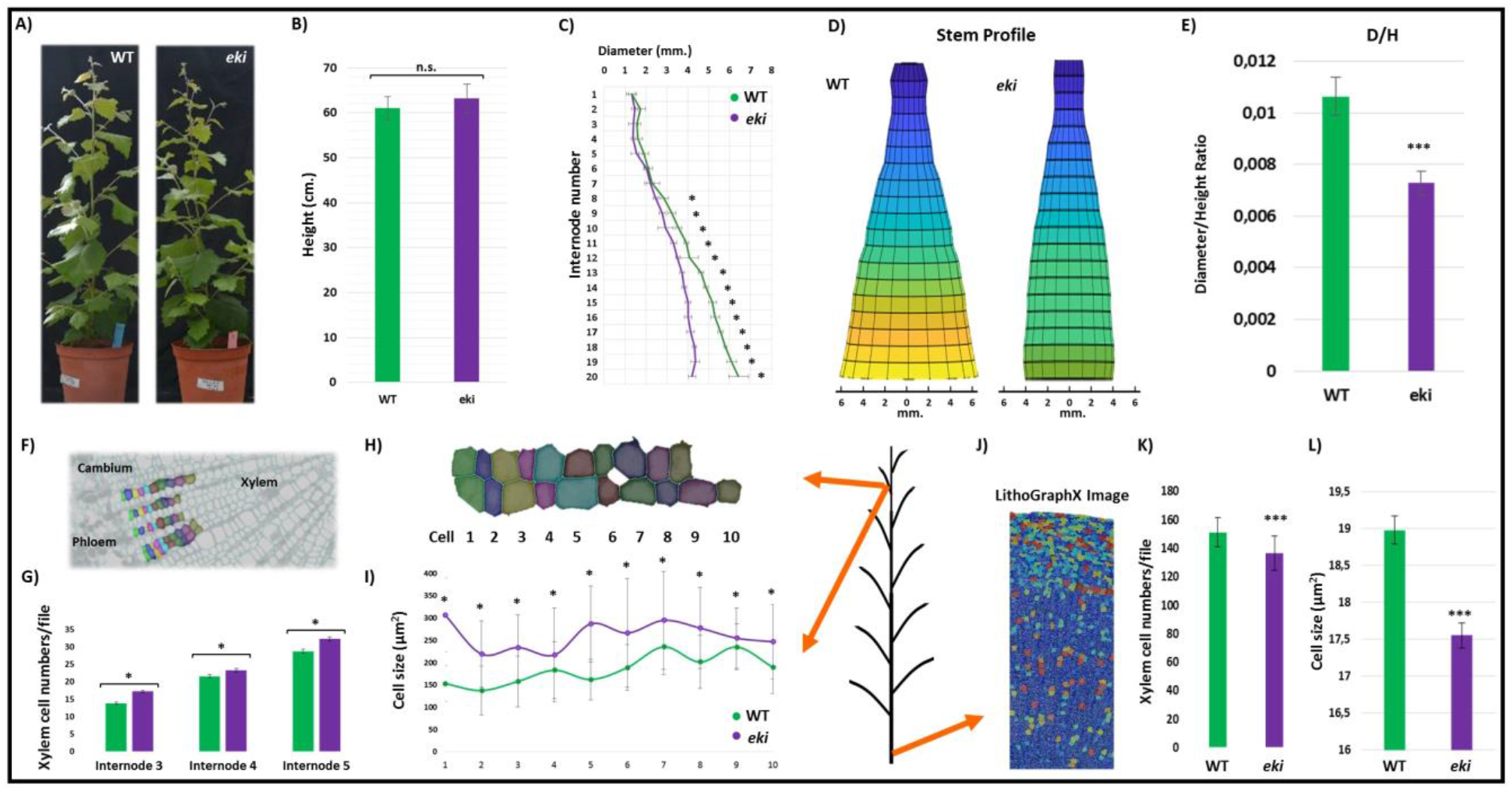
The *eki* phenotype increases cell size and numbers in young internodes but restrict them at the stem base. A) Phenotypes of clonally propagated WT and *eki* BC_1_ trees, grown for 3 months. B-C) Stem allometry showing total height and the diameter of each internode. D) Stem profile reconstructed from data in C highlights the decrease in the thickening slope of the *eki* trees. E) Diameter (in the basal internode) vs. total height ratio. F-I) Early stages of xylem expansion in top internodes from young trees (one month after planting). G) Xylem cell numbers per individual xylem cell file on top internodes. H) Segmented progression of most recent 10 xylem fibers cells developed from cambium. I) Cell sizes (µm^2^) across the developmental progression in F, H. J) Image segmentation of stem base by LithoGraphX (color code = cell size in µm^2^). K) Xylem cell numbers per individual xylem cell file counted in the stem base. L) Cell size from randomly sampled xylem fiber cells at the stem base. The schematic figure indicates the analyzed stem positions. Asterisks indicate statistically significant differences \**p* ≤0.05, \*\*\**p* ≤0.001 (Student t-test), by comparing WT to *eki*.

The same experimental setup described above for WT trees was simultaneously used for clonally propagated BC_1_ *eki* trees. The results showed that, in contrast to the WT response, in *eki* (T4/T1) the diameter increments of basal internodes were similar in all scenarios (**Fig 1B**). The same pattern was shown when upper internodes were included in the analysis (**Fig S1**). The pulling scenario slightly increased the average radial growth but this difference was not statistically significant. Unlike other scenarios, pulling also reduced the data dispersion in *eki* (C_V_=0.065) to WT (scenario 1) levels (C_V_=0.069) (**Supplementary Data 2**), suggesting that trees grew more homogeneously than in other scenarios. In addition, the response patterns comparing weight manipulations vs. no weight were different to those observed in WT (**Fig 1D**). Here the variations across time points or stem positions were very small, except for the scenario 2 (added weight on supported trees) in which response pattern seemed more similar to those in WT scenario 3 and 4.

Combined, these results indicate that before the collapse *eki* cannot adjust its growth proportions and respond to vertical changes of the stem weight, as trees were resistant to the mechanical treatments. The reduced radial growth after three months, further led to a stem mechanical failure that was manifested in the collapse of *eki* trees.

### The *eki* secondary growth phenotype is differentially manifested along the stem

The later manifestation of the radial growth phenotypes and the missing response to stem loadings, indicates that unlike WT, the *eki* trees can not compensate by radial growth for the increasing tree size. The reconstruction of diameter profiles across stem internodes shows that while in WT every internode is thicker than the previous one (from top to bottom), in *eki* the slope of the growth curve is strongly reduced (**Fig 3D**). Furthermore the diameter of the *eki* main stem is significantly reduced starting from the internode 7 (**Fig 3C**).

To dissect the anatomical phenotypes of xylem and phloem tissues, we quantified cell numbers and sizes in progressive developmental stages along the stem. In three month old trees the youngest internodes of *eki* underwent xylem expansion earlier than WT trees (**Fig S3E**), also one month old trees showed increased numbers of xylem cells (**Fig 3G, S3F**). The analysis of segmented images provided an accurate characterization of xylem cells by comparing cell files (of fibers) within the radial developmental progression in WT vs. *eki* (**Fig 3F, H**). This approach revealed that xylem fiber areas were larger in *eki* as compared to WT (**Fig 3I**). On the contrary, the oldest internodes at the base showed the opposite phenotype. Xylem tissues had significantly less xylem cells per file (in average 15±4) (**Fig 3K**), but the area occupied by individual xylem fibers was smaller in *eki* as compared to WT (**Fig 3J, L**). Therefore, the xylem cell sizes, and not the cell numbers, contribute to the primary reason for the impaired secondary growth phenotypes in *eki*.

The radial anatomy analyses indicated that the xylem phenotypes in *eki* depend on the height or developmental stages (size) of stem internodes. The *eki* phenotype promotes xylem cell expansion and proliferation in young internodes, but reduces cell numbers and most importantly cell sizes in the older internodes at the base. The last phenotype corresponds to the bending point during the *eki* collapse.

### Secondary growth creates radial stress patterns in the stem

Plant morphogenesis inherently affects the mechanical environment that surround the developing tissues. This process is the result of compressive and tensile forces produced by cell proliferation and cell expansion. During secondary growth the cambial activity presumably creates mechanical conflicts by pushing tissues inwards during xylem development. However mechanical stress patterns derived from secondary growth have so far not been visualized in stems or roots cross-sections. Our previous observations indicate that in *eki*, xylem size and probably cell expansion are affected in a context dependent manner, relative to the stem developmental stages or height of the stem. Thus, cell size may indirectly reveal whether a particular context promotes or restricts cell expansion. Therefore, we studied the radial pattern of cell expansion at different stem heights (**Fig 4**).

**Figure 4.**
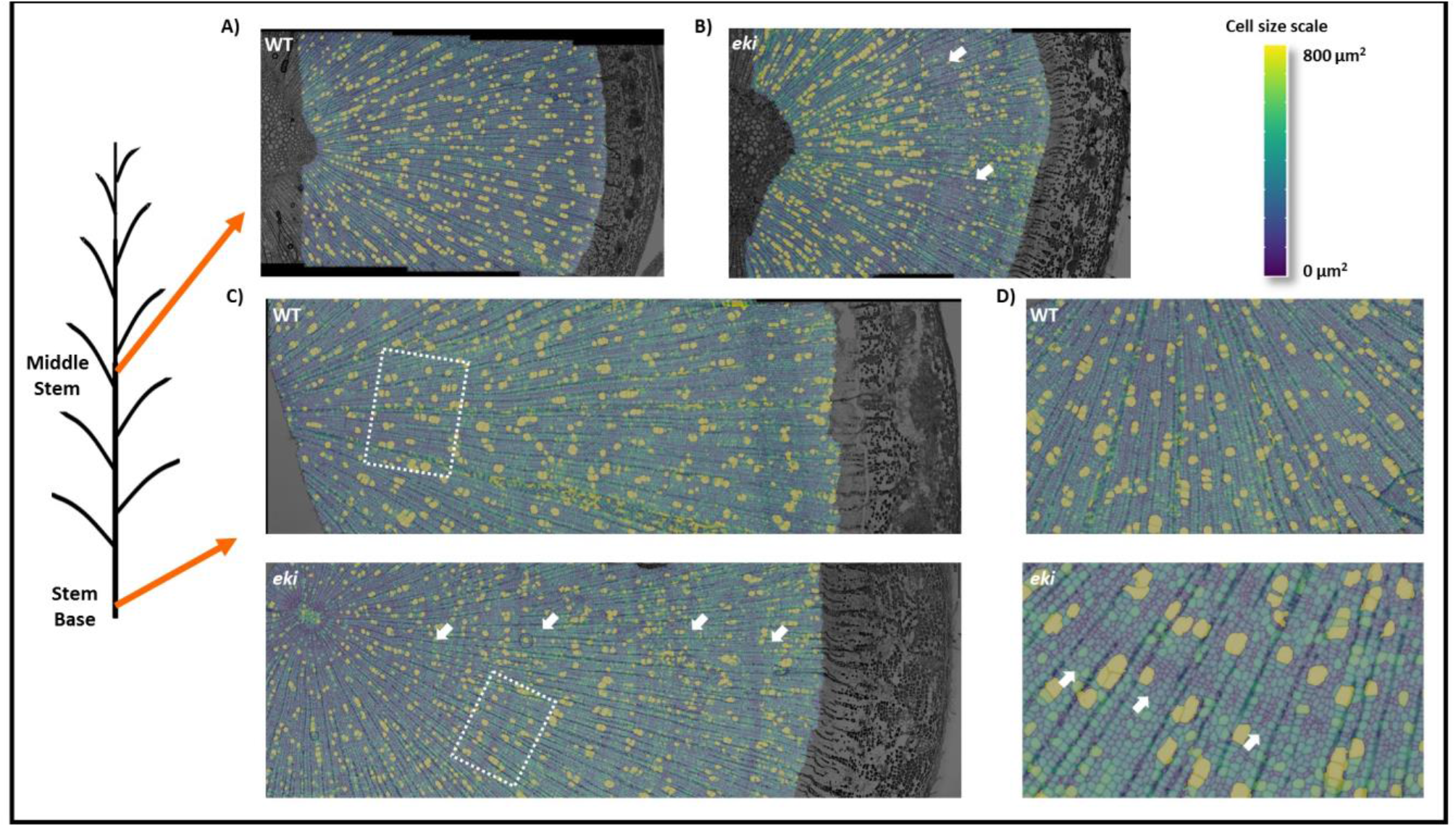
Growth stress patterns in *eki* stem correspond to defects in cell expansion. Anatomical images were segmented with LithoGraphX and overlaid with heatmaps corresponding to cell size (µm^2^). Schematic figure indicates the analyzed stem positions. A) WT and B) *eki*, stem cross-section taken at middle stem height. White arrows indicate concentric rings of small cells. C) WT and *eki* stem base cross-sections. D) Magnified images from C). Stress patterns occurred in 0/5 in WT and 5/5 in *eki*.

LithoGraphX segmented images of the stem cross-sections, showed the distribution of individual cell sizes. The internodes at the middle stem height, showed in WT a heterogeneous distribution of cell sizes with larger individual cell areas for vessels, and smaller ones assigned to fibers. This heterogeneous pattern was maintained at the from the center of the stem (pith), to the cambium (**Fig 4A**).The *eki* stems at the same height had sectors with reduced fiber cell size, but interestingly, they occurred only on the cambium side of the cross-section and not near the pith (**Fig 4B**). To understand if this phenotype is also developmentally associated we studied internodes at the base of the stem. In this section WT trees had a similar pattern to that described before (**Fig 4C, D**). Contrastingly, in *eki* trees the pattern fiber cell size reveals sectors of smaller cells arranged in concentric rings. These sectors were found even near the center of the stem (pith) (**Fig 4C, D**).

In stems, the developmental progression can be traced back in time in the longitudinal direction. Given younger internodes of *eki* displayed only a partial phenotype near the cambium (most recently developed xylem), and the pattern was almost complete (from cambium to pith) in the older internodes at the base, these observations indicate that the cell size reduction can also occur later, when fibers are fully developed. In addition, the fiber size phenotype of *eki* is not randomly distributed but follows an intermittent radial pattern, which may highlight the lateral constraints during the radial expansion of secondary growth.

### Cell wall integrity is affected in *eki* trees

The structural failure in the main stem, and the phenotypes associated with cell expansion could be partially explained by defects in the cell walls composition or structure. To investigate this aspect we performed a global profiling analysis of cell wall composition by separating bark and wood samples, from young internodes (IN:6-8) and mature ones (IN 9-11). On this set of samples we studied the cellulose, hemicelluloses and lignin. The results from lignin composition identified a slight increase of total lignin in *eki* only in young bark samples (+8.6%). In *eki*, G-type lignin was increased in the bark and S-type lignin decreased in the xylem (**Fig. S6**). The composition of hemicelluloses was assessed by measuring the content of their building monosaccharides (**Fig. S7**). In this analysis, young internodes from *eki* had an increase of xylose and non-cellulosic glucose in the bark, and arabinose, fucose and mannose in xylem. Mature internodes in *eki* had slight increase of non-cellulosic glucose in the bark and arabinose in the xylem, the remaining monosaccharides showed no significant differences. Finally, no detectable differences were found in total cellulose in any tissues (**Fig. S8**).

Altogether, the differences of cell wall components were more evident the young than in the old internodes. However, given the unspecific nature of these differences and the fact that changes were relatively subtle, the cell wall profiles in *eki* seem to represent complex defects associated to the development or integrity of cell walls, rather than the defect in one main component.

### Patterning of secondary cell wall deposition is affected in *eki* trees

While the final cell wall integrity was affected in *eki*, our previous anatomical analysis revealed morphological differences in the wood forming tissues. Therefore, we further investigated whether these patterns correlate with the cell wall composition *in situ*. Crystalline cellulose was visualized through CBM3a immunolocalization (**Fig. 5A**). In the WT sections the immunolabeling pattern showed that cellulose was evenly distributed in xylem, starting from the very first developing xylem cells. On the contrary, the same sector showed an irregular pattern in *eki*. Lignification was visualized by UV autofluorescence over thin cross-sections (**Fig. 5B**). Images acquired with the same exposure time revealed that the lignin autofluorescence increased rapidly in developing xylem of WT stems; the same tissues revealed a darker area in *eki* which showing a delay in the lignification of secondary xylem tissues.

**Figure 5.**
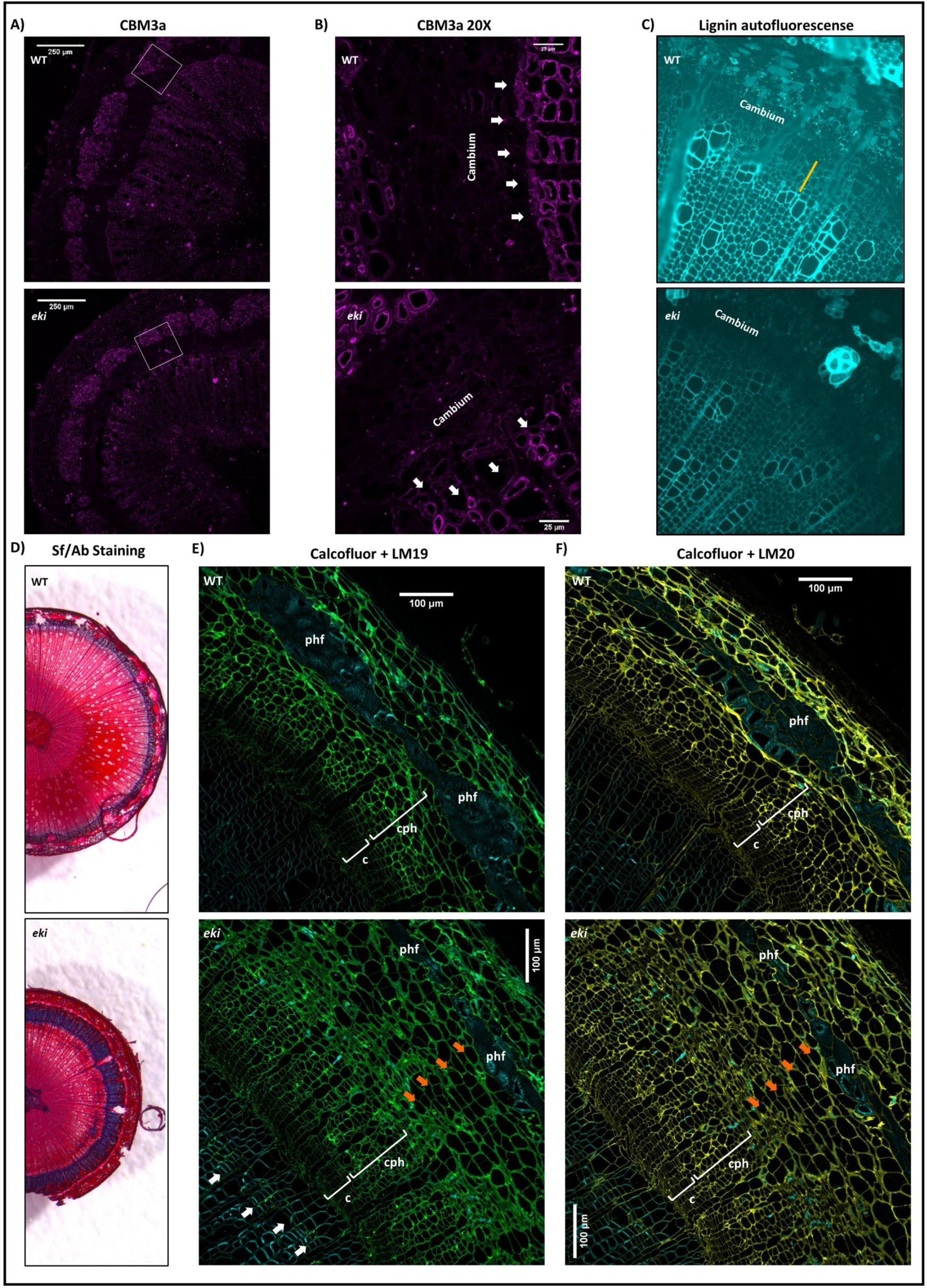
Patterning of cell wall polymers in the wood forming zone. **A)** Immunolocalization of crystalline cellulose by CBM3a, and **(B)** magnified image of cambium and developing xylem (right panel). White arrows indicate the first developing xylem fibers **C)** Visualization of lignin by UV autofluorescence. Orange line indicated non-lignified developing xylem fibers. **D)** WT and *eki* cryosections from the base of the stem, stained with alcian blue and safranin. **E-F)** Immunolocalization in the oldest internode at the stem base. Images correspond to the calcofluor white counterstaining (cyan), and **E)** LM 19 (green) or **F)** LM 20 (yellow) immunolocalization. Brackets in the figures indicate the cambium (c), conductive phloem (cph). Phloem fibers (phf). White arrows indicate the LM19 labeling in xylem. In WT trees conductive phloem connects cambium and phloem fibers, but extra layers of parenchymatous cells labeled by LM19 and LM20 can be found in *eki* (orange arrows).

Next, we screened for further phenotypes from histological cryosections. Unlike other methods, the cryosections are cut from fresh samples and require no additional treatments other than staining. The histology of fresh tissues showed that WT cross-sections had only a narrow alcian blue stained sector within the cambium zone (**Fig. 5C**). This cationic dye preferably stains acidic and pectin-like polysaccharides typical for primary cell walls (23). The same staining showed a broader area in the *eki* stem that expanded from the phloem, over the cambium and across the developing xylem (**Fig. 5D**). To verify if this pattern corresponds with pectin polysaccharides we performed immunolocalization using LM 19 and LM20 antibodies that label specifically homogalacturonan in its demethyl-esterified (LM19) and methyl-esterified (LM20) form. We then studied cross-sections from three stem positions: top (IN5), middle (IN12), and base (**Fig 5E-F, Fig S9**). Both antibodies showed preference for cambium and bark areas, as these tissues are rich in primary cell walls. The anatomy of WT cross-sections showed the developmental progression of cambium, conductive phloem and phloem fibers, in which the first two tissues are strongly labeled by both antibodies. Instead, *eki* cross-sections showed LM19 and LM20 labelling in parenchymatous cell types, which were positioned between the conductive phloem and phloem fibers (**Fig 5E-F**). Moreover, the development of the phloem fibers was strongly restricted in *eki*. Finally traces of LM19 labeling was also found in xylem tissues of *eki* samples, but not in WT. Altogether these analyses indicate that the development of cell walls was affected and delayed near the meristematic zone and towards mature tissues. Furthermore, the enlarged sectors in *eki* stained by Alcian blue and detected by LM19 and LM20, suggest that cell walls in developing phloem and xylem tissues of *eki*, retain characteristics of primary cell walls (being rich in pectin) even outside the cambial zone.

### Tissue-specific mapping of elasticity reveals a different mechanical environment around the cambium of *eki* trees

The morphological changes in the anatomy and defects in cell walls might have an impact on the mechanical properties of the stem. Therefore, to dissect the nature of the physical failure in *eki* trees we performed series of mechanical tests at the tissue and organ level.

First, we aimed at producing a map of elastic modulus at the tissue level across wood forming zone by performing indentations with a 25µN bead AFM tip on fresh stem cross-sections (**Fig 6**). To provide a perspective of the developmental progression, we expanded the AFM mapping across the phloem, cambium, and xylem. In cross-sections from WT trees, the elasticity maps revealed areas with reduced Young modulus (soft areas) within the cambium and phloem tissues (**Fig 6A**). On the opposite direction, developing xylem showed a pattern of increasing Young modulus (stiffer) which eventually becomes very compact. In agreement with these observations, the elasticity histograms shift from a distribution with majority of soft measurements (in phloem and cambium) to a nearly normal distribution (in xylem). On the other hand, the elasticity pattern was different in *eki* cross sections (**Fig 6B**). Cambium and phloem tissues had a low Young modulus (softer), and the developing xylem in *eki* showed areas of low Young modulus, that only later revealed stiffer spots similar to those of mature xylem in WT.

**Figure 6.**
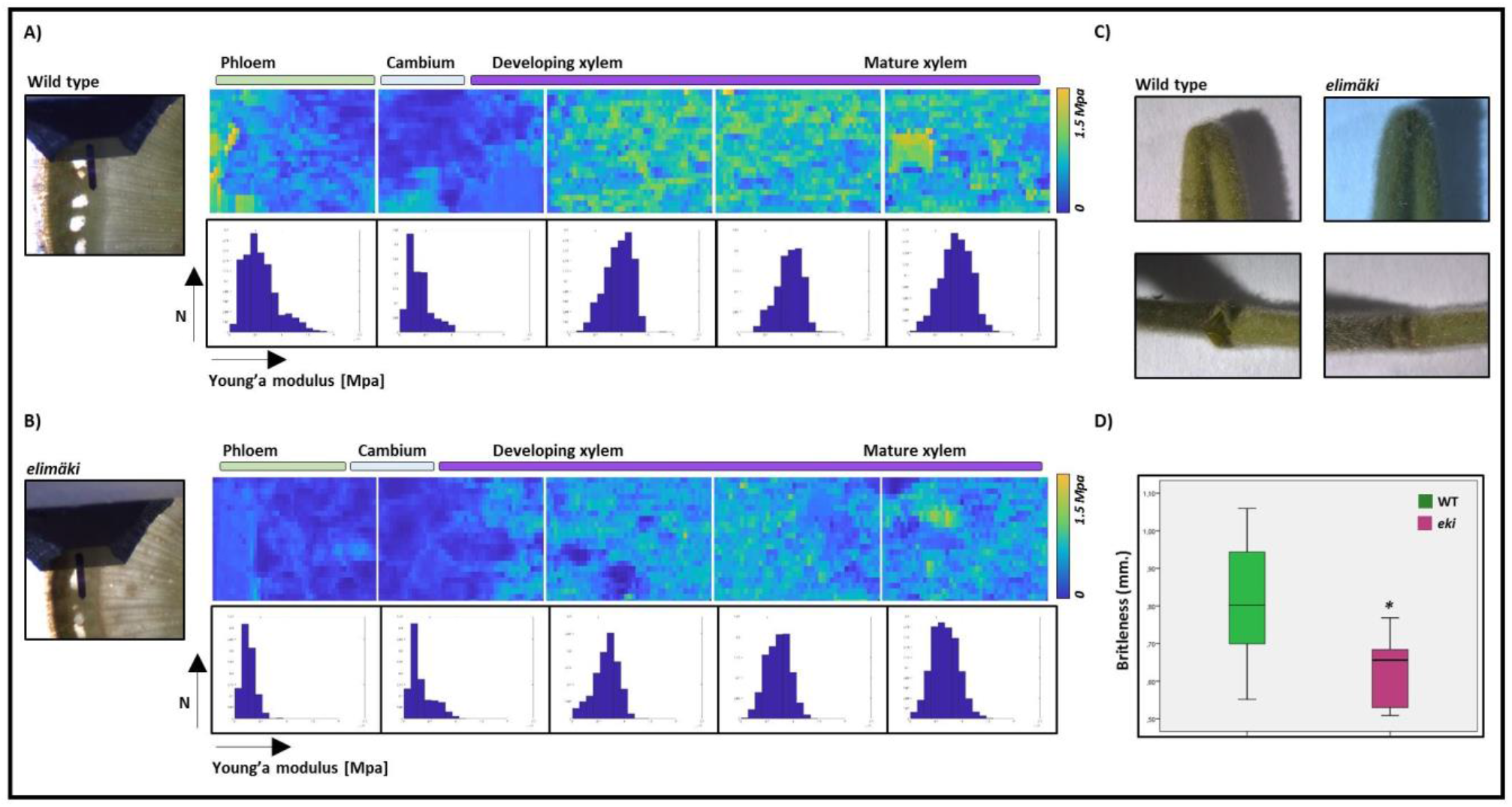
Tissue-specific mechanical properties across the wood forming zone. AFM based reconstruction of Young’s modulus across phloem, cambium and xylem tissues in WT (A) and *eki* (B), in stem cross-sections from middle stem (IN12). A stereoscope image was acquired before each run to identify the tissues in the sample (image shown corresponds to the first map). Heatmaps shown in the figure are the representative maps of 13 WT and 14 *eki* analyzed stem cross-sections in two independent repetitions. Color scale from 0 to 1.5 Mpa. Histograms in the lower panels illustrate the distribution of Young’s modulus measurements for each map. C) Outcome of manual bending of fresh stem internodes of approximately same diameter. D) Results from three-points bending test showing the brittleness in air dried stem internodes of approximately same diameter (n = 9).(*) Asterisk indicate statistically significant differences *p*≤0.05 (Student-t test), by comparing WT vs. *eki*.

On a larger scale, when fresh stems were manually bent the WT internodes broke on the bark, whereas the *eki* sections did not break and appeared to be more flexible (**Fig 6C**). Therefore, to assess the lateral tensile strength of complete internodes in controlled conditions we dehydrated stem sections, and performed three-point bending test (**Fig 6D**). In this assay *eki* internodes were significantly more brittle than in the WT. In addition, the hardness of WT samples shared a high positive correlation with the diameter of internodes, but this correlation was lost in *eki*. Combined these results indicate that the mechanical properties of developing tissues in *eki*, are already modified in the vicinity of cambium making them softer. Later, the overall structural integrity of stem internodes is compromised.

### The transcriptome of *eki* overlaps with the *Arabidopsis* touch response

The earliest anatomical phenotypes in the stem of *eki* appear young internodes, represented by proliferation of xylem cells in internodes 3-4-5. Therefore, to study the transcriptional profiles we dissected the previous internode with no morphological differences, internode 2 (IN2) and the internode 5 (IN5) where the phenotype was very clear in both WT and *eki*. Since this internodes represent the earliest phenotype progression, we also dissected the IN5 in bark and wood. More than 300 million raw reads pairs were produced from four biological replicate libraries for each sample (**Supplementary Data 3**). After data trimming and quality control, an average of about 71 million high quality clean reads were generated per sample, with an average overall alignment rate of 92.5% to the *B. pendula* genome. The average mapping rate of reads mapped to the coding regions of genome was 71.45%. PCA results showed good correlations between the biological replicates of each sample (Figure. S8). Comparative analysis of the relative levels in gene expression identified 846, 1004 and 1034 differentially expressed genes (DEGs) in the comparison pairs of IN2, IN5-bark and IN5-wood between *eki* and WT, respectively (**Supplementary Data 3**).The global changes in WT vs. *eki* gene expression were captured by the GO enrichment analysis of DEGs (**Fig. S10**). The GO categories associated to cellular component highlighted for both IN2 and IN5-bark “intrinsic component of mitochondrial membrane”, whereas the same categories correspond with “plant-type cell wall” in IN5-wood. Concerning the categories associated to biological processes the top enriched terms were “flavonoid biosynthesis” in IN2, cellular response to nitrogen starvation in IN5-bark, and “respiratory burst” and “jasmonic acid biosynthesis” in IN5- wood. Overall this set of enriched GO terms corresponds with abiotic stress, but also the very same molecular processes such as mitochondrial stress, production of reactive oxygen species and jasmonic acid signalling have been directly associated with the mechanosensing response of *Arabidopsis* to touch (thigmomorphogenesis) (24–26). To test if our list shares such a similarity, we compared the DEG in WT vs. *eki* with DEGs after touch induction in *Arabidopsis* (25). Interestingly, 12.3% of *Arabidopsis* DEGs upon affected touch induction are also differentially regulated in WT vs. *eki* (**Fig. 7**). Touch inducible genes such as WRKY15 and WRKY40 were upregulated in *eki* samples, in both IN5 bark and wood; whereas OUTER MEMBRANE PROTEIN 66 was downregulated. In addition, 6 genes belonging to the Mechanosensitive ion channel family (MSL) were differentially expressed, two upregulated and four downregulated in *eki* IN5-bark. The mitochondrial stress marker, *Alternative Oxidase 1a* (AOXa) was significantly downregulated in *eki* IN2 and IN5-bark. The jasmonic acid biosynthesis gene AOS is required for touch induced developmental responses in *Arabidopsis* (24), this gene was also significantly downregulated in *eki* IN5-bark and wood. Another gene required for thigmomorphogenesis is VIP3 (27), this gene was significantly upregulated in *eki* bark. The birch orthologous genes of *Arabidopsis* TOUCH3 and TOUCH4 had barely detectable expression in our samples. The results from the GO enrichment analysis suggest that the differences in WT vs *eki* transcriptomes are associated to mitochondrial stress, and later in the IN5-wood samples with cell walls; which corresponds with the xylem phenotypes. Interestingly, we also found that the expression of genes associated with thigmomorphogenesis and mechanosensing is affected in *eki*. Given that stem samples were collected from untouched trees, this molecular pattern suggest a constant transcriptional background in *eki* that interacts in different directions (upregulating or downregulating genes) with the complex mechanosensing pathway.

**Figure 7.**
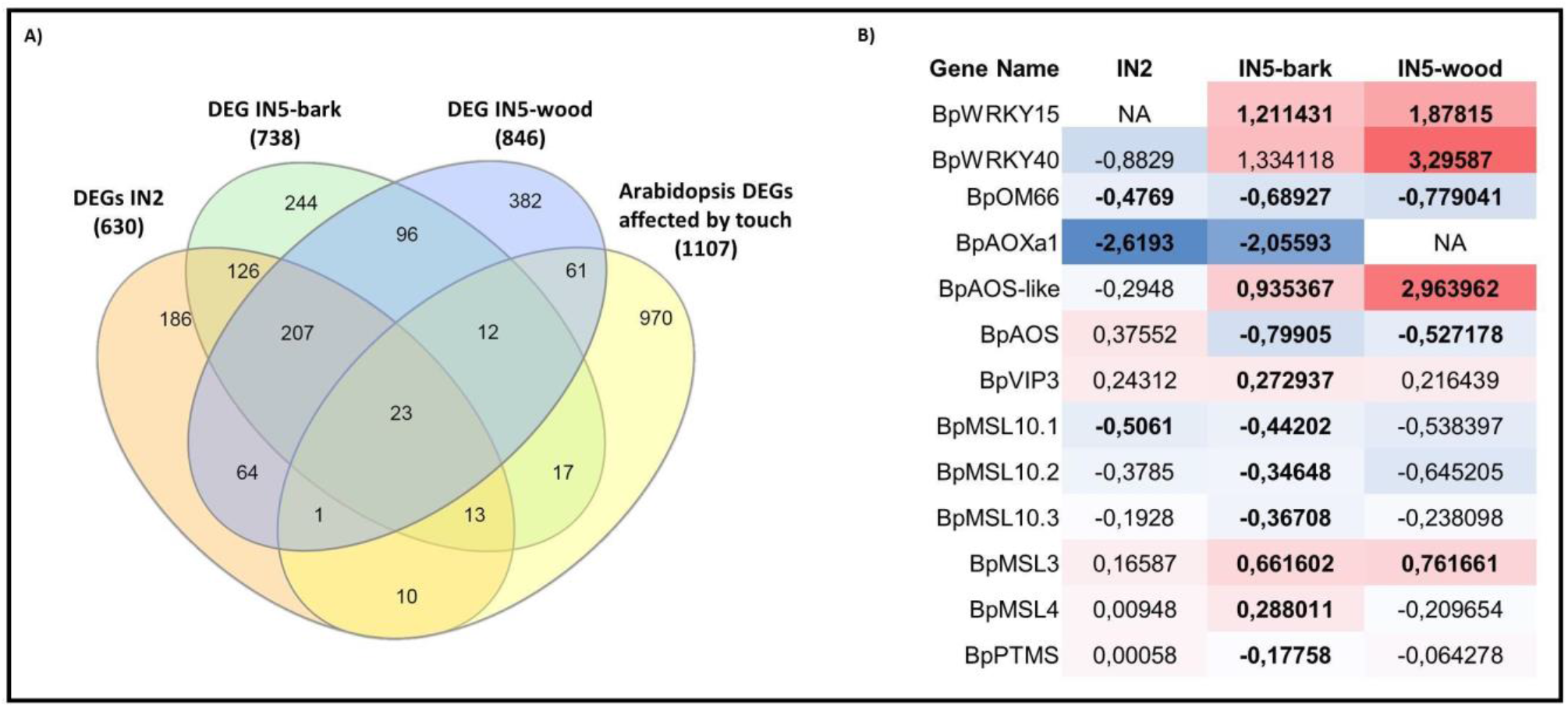
The DEGs in WT vs. eki trees shares similarities with the Arabidopsis thigmomorphogenesis response. **A)** Venn diagram of Arabidopsis orthologues of birch DEGs in IN2, IN5-bark, IN5-wood compared to the list of DEGs induced by touch in Arabidopsis (25). **B)** Candidate genes involved or required for thigmomorphogenesis in Arabidopsis and their corresponding birch orthologue. Values represent Log_2_FC in WT vs. *eki* (negative value = downregulated in *eki*), in bold numbers *p* adj ≤ 0.05.

## Discussion

Trees do not belong to a monophyletic group, instead the distinctive features of tree-like plants such as secondary growth and woodiness, evolved multiple times during the diversification of Embryophytes (28, 29). Such evolutionary history suggest that growing thick and tall was positively selected multiple times in the context of similar survival and reproductive demands. Lifting the biomass is therefore a physical challenge common to all tree species. The additive effect of growth, increases the weight and physical stress, which in turn regulate the localization and magnitude of growth. This feedback loop, resembles the perception of tissue deformation during gravitropism which in turns modulates the straightening process of stems, this mechanism was also referred to as “proprioception” (2). However, in this work we studied the components of proprioceptive responses in a different mechanical context of stems. Standing trees differ from tilted or bent stems in orientation, type of mechanical conflicts and consequently in developmental responses. Here, the weight vertically loads on the stem column in parallel to the gravity vector. Simultaneously, the radial growth increases compressive forces in the wood forming zone, and accumulates tensile stress at the periphery (14). While some of these forces have been measured by interpreting the residual stress (30), the extent to which this mechanical conflicts surrounding the cambium can conditionate wood formation remains hypothetical.

In our experiments with vertical weight manipulations we detected differences of height and diameter within 3 weeks. Only semi-free trees with extra weight added underwent significantly more radial growth compared to other treatments. These results indicate that slight movements of stem posture are important to perceive and respond to additive loadings in the stem. Since stem movements normally induce tissue deformation from lateral tensile and compressive forces, proprioception of standing trees may share mechanically some aspects with gravitropism. At the same time, the response of stems was also promoted in upper internodes indicating that the radial growth response is systemic. While here our experimental design detected the first response of stems, further studies with longer treatments and additional conditions (e.g. different weights) are required to further model the long-term responses involved in proprioception.

The identification of a mutant birch defective in proprioceptive responses, and the Mendelian segregation of this trait, enabled an insightful comparison of WT vs *eki*. Although forward genetics with trees have been reported before (31–33), this process normally takes decades. This is the first time a BC_1_ segregating population is obtained in 4 years. In agreement with the trait segregation, the QTL mapping identified one locus (EKI) linked the *eki* phenotype and 76 DEGs within this mapping window. As revealed by the Marey plots, the target locus was located in a chromosomal region with low recombination events compared to the other chromosomes (**Fig S3B**). Therefore, larger population numbers would be required to identify the causative mutation.

The characterization of *eki* uncovered a complex phenotype in which younger internodes had increased xylem cell size and numbers, but older internodes at the base displayed the opposite. This indicates that xylem phenotypes in *eki* are inversely associated to the size, weight or developmental stage of the samples.

In addition the cell wall integrity and elasticity of developing xylem cells was compromised in *eki* making them softer as indicated by the AFM measurements. Therefore, one hypothesis is that a more elastic mechanical environment in younger internodes promotes cell proliferation and expansion, however weight and growth-derived mechanical stress restricts this process in older internodes at the base. An analogous effect from the mechanical environment on the developmental progression of a plant meristem has been reported in *Arabidopsis* (16). In line with this, we identified rings patterns in *eki* stems indicating sectors with reduced cells size which may result from the mechanical stress produced by radial growth and the lack of rigidity in *eki* cell walls. Furthermore, similarly to previous the previous phenotypes the growth-stress rings were more frequently found in older internodes indicating that a larger tree size inversely correlates with the strength of the phenotypes.

At the transcriptional level the analysis of DEGs in *eki* indicated the many misregulated genes were part of mitochondrial stress pathways. This transcriptional profile is also triggered in the *Arabidopsis* mechanosensitive response to touch (25, 26, 34). Therefore, while the primary cause of the *eki* phenotype is far from being resolved, the ELIMÄKI locus seems to be required for controlling the retrograde signaling pathway. We speculate that the ubiquitous activation of this stress condition masks the proprioception capacity of *eki* stems. When *Arabidopsis* stems are mechanically stimulated stem elongation is restricted in response. Interestingly, the screening for *Arabidopsis* mutants resistant to touch resulted in very different candidate genes such as jasmonic acid biosynthesis enzymes (24), or transcription associated proteins (27), and these genes were also affected in *eki* highlighting the fact that plant mechanosensing is a complex phenomenon with multiple pathways being involved (35).

Altogether, we conclude that the proprioceptive response of standing trees can be triggered by weight as long as stems can move, but also this mechanism is intrinsically conditioned by the composition and elasticity of cell walls in the wood forming zone. Finally, using birch as a model organism, this forward genetics approach could be a powerful tool in basic and applied research of trees species.

## Acknowledgements

The authors are grateful for the excellent technical assistance of Katja Kainulainen, Wouter Beijk and Rebecca Kramps. This study was supported by Finnish Centre of Excellence in Molecular Biology of Primary Producers, Academy of Finland (CoE 2014-2019) (271832), the Gatsby Foundation (GAT3395/PR3), University of Helsinki (799992091), and the European Research Council Advanced Investigator Grant SYMDEV (323052).

## Authors Contribution

Y.H, K.N and J.A.S, designed the research; J.A.S, X.S, A.P, P.R, M.B, J.I, J.T, H.K, G.E, S.M, H.H-R-R, O-P.S, C.S, R.H, A-P.M and K.N performed the research; J.A.S, X.S, A.P, P.R, M.B, J.I, J.T, H.K, G.E, O-P.S, L.G, O.S, J.S and K.N analyzed the data; and J.A.S, Y.H and K.N wrote the paper.

## Methods

### Induced early flowering and crossings

The natural flowering period for birches in southern Finland is spring (April to May) when trees break dormancy. Therefore, in the first year (2011) the identified cultivar *Betula pubescens cv. Elimäki* (EO) was selfed to produce S1 trees by performing controlled crossings. Briefly, branches with dormant male flowers were bagged with pollination bags (model: 2D-1-1W, from PBS International, UK) during March 2011. Within the following weeks male flowers burst, and female flowers emerged inside the closed bags producing S1 seeds. F1 trees were obtained by collecting wind-pollinated seeds from EO (July-August 2012). All seeds were allowed to ripen in the tree, air dried for 5 days and stored in −5°C freezer for at least one month. In March 2012, seeds were planted and trees were grown in normal greenhouse conditions (18hrs DL) put in dormancy (October 2012) by allowing greenhouse rooms to acclimate to outdoor conditions. In total 21/21 F1 trees had WT phenotype, and 15/15 S1 trees had mutant phenotype. Flowering of F1 trees was induced in February 2013 when there was no registered ambient pollen, by using the previously published protocol (19). F1 trees flowering was successfully induced (2013), but this seeds didn’t germinate. On the second year of F1 trees flowering was successfully induced again (February 2014), producing viable seeds. After visible female flowers emerged, branches were bagged and pollinated by injecting EO pollen. Seeds were allowed to ripen in the branch, air dried and stored in −5°C freezer for at least one month. The BC_1_ population was then grown from seeds in February 2015. The BC_1_ population resulted in 60 trees with 33 WT phenotype and 27 *eki* phenotype.

### Plant material and growth conditions

The cultivar *Betula pubescens cv. Elimäki* (EO) is located in DD: 60.751247, 26.489245 and was identified by Natural Resources Institute Finland (Luke ID: E11431). Individuals 14-038-01 (*eki*), 14- 038-22 (*eki*), 14-038-58 (WT), 14-038-17 (WT), were established *in vitro* and clonally propagated as previously described (36) and grown in normal greenhouse conditions (22-25°C,18hrs light) with automatic watering. Unless otherwise indicated in the figure legends the analysis of trees was performed in 3 months old trees or before collapsing of mutant individuals.

### Vertical mechanical manipulations

The vertical mechanical manipulations consisted in 4 different experimental setups. Weight treatments were performed by manipulating the equivalent to 75% of the weight in upper half of the stems. This weight was estimated by cutting and measuring such stem sections in 3 WT and 3 *eki* trees weight, and then using this information to calculate the respective 75% weight in WT or *eki*. Weight was applied by using handmade rings of modelling clay (Staedtler). Then, weight treatments were distributed in a gradient (heavier rings in the bottom, and lighter in the top) positioned at 50%, 62.5% and 75% of stem height, the weight of each ring also mimics the biomass weight distribution or loading gradient in WT and *eki* trees.

Scenario 1: Trees were supported by three sticks next to the stem to prevent movements, and no weight manipulations were performed. Scenario 2: Trees were prepared as Scenario 1, but weight was applied as described. Scenario 3: Only one stick was added 12cm. away from the stem; in the upper tip of the stick we added a metal wire ring (D=25cm.). Next weight was also applied as described. This Scenario permits stem oscillations which are prevented in Scenario 1 and 2. Scenario 4: Pulling was performed using fishing lines (Sufix 832). Lines were attached to the stem with a tick protective parafilm layer. A metal structure was built on the top of trees (1.5 M) to allow hanging of Eppendorf tubes filled with weight (fishing sinkers). The weight amount and distribution was exactly the same as that calculated before for weight rings.

### Anatomy analysis

Samples used in cell number and cell size quantifications were fixed (1% glutaraldehyde, 4% formaldehyde, 0.05M sodium phosphate, and after dehydration, embedded in Leica HistoResin. 5 µm thin cross-sections were made in Leica RM2165 rotary microtome, with a Leica microtome knife. Afterwards, slides were stained in a 0.05% toluidine blue solution, and imaged using a Leica 2500 microscope. Image analysis was done in Fiji (37) and cells segmentation and size quantification was done in LithoGraphX with Builder 1.2.2.7 (38, 39).

Transition Electron Microscopy (TEM) of tension wood samples was performed as previously described (40) without staining. Samples were processed and imaged at the Electron Microscopy Unit, Institute of Biotechnology University of Helsinki.

### Pyrolysis-gas chromatography and mass spectrometry (Py-GC/MS)

50 µg (± 10 µg) of ball-milled wood and bark powder was applied to a pyrolyzer equipped with an auto sampler (PY-2020iD and AS-1020E, Frontier Lab, Japan) connected to a GC/MS (7890A/5975C; Agilent Technologies, Santa Clara, CA, USA). The pyrolysate was separated and analyzed according to (41).

### Monosaccharides and cellulose composition analysis of cell walls

Alcohol insoluble residue (AIR) was prepared from fine wood and bark powder by extraction with 80% ethanol for 30 min at 95°C, repeated with 70% ethanol, followed by chloroform:methanol (1:1) and two washes with acetone at room temperature. In order to remove starch, the powder was treated overnight at 37°C twice, with α-amylase from pig pancreas (Roche 10102814001; 100 units per 100 mg of AIR) in 0.1M potassium phosphate buffer, pH 7.0. Monosaccharide composition was analyzed using trimethylsilyl (TMS) derivatization method. 500 μg of amylase-treated material and 30 μg of inositol, used as internal standard, were methanolysed by 2 M HCl/MeOH at 85°C for 24hrs. Derivatization was carried out using Tri-sil reagent (3-3039 Sylon HTP kit, Supelco, Sigma-Aldrich) at 80°C for 20 min as previously described (42). The silylated monosaccharides were determined on a GC/MS (7890A/5975C; Agilent Technologies, Santa Clara, CA, USA) according to (43), using a J&W DB-5 MS column (Agilent Technologies) with the oven program: 80°C followed by a temperature increase of 20°C/min to 140°C for 2 min, then 2°C /min to 200°C for 5 min, then 30°C /min to 250°C for 5 min. The total run time was 47 min. Crystalline cellulose content was determined using the Updegraff method (44) followed by an anthrone assay to detect the released glucose (45).

### Cell wall polymers immunolocalization

Trees sections of WT and *eki* birch were dissected with razor blades (several mm thick) and were immediately fixed for in a solution consisting of 4% formaldehyde (freshly prepared from paraformaldehyde powder, Sigma) and 0,5% glutaraldehyde (Sigma) in a 0,1M phosphate buffer pH7. Fixation, dehydration and resin infiltration steps were all micro-wave (MW) assisted using a PELCO BioWave Pro (Ted Pella, Redding, CA). Fixation was realized at 150W, under vacuum (20Hg) (5x 1’). Samples were left in the fixative overnight at 4°C and then washed 3 times in PBS. Samples were then processed through increasing dehydration steps (25%, 50%, 70%, 90%, 96%, 3x 100% Ethanol, vacuum 20Hg, MW 150W 5’). Resin infiltration (LR White medium grade, Agar scientific) was then realized through increasing resin concentration: 33% Resin in ethanol 100%, 66% Resin in ethanol 100%, and 3 times 100% Resin (20Hg, MW 200W 5’). Samples were left overnight in 100% for effective resin penetration in the samples. Resin polymerization was subsequently realized at 60°C during 17h. Semi-thin sections (1µm) were then obtained with a Leica EM UC7 ultramicrotome. LM19 and LM20 Immunolocalization on semi-thin sections was performed as follows: blocking step (BSA 2% in PBS, 1mL per slide, 1h RT); primary antibodies 1/10 in BSA 2% in PBS, 500µL per slide, ON 4°C (LM19/LM20 (46), plantprobes, Leeds UK); 5 washes in BSA 2% in PBS (MW 150W, 1’); secondary antibodies (Alexa Fluor 488 Goat anti-Rat IgM, ThermoFisher Scientific, A-21212) 1/100 in BSA 2% in PBS, 500µL per slide, 2h RT; 5 washes in BSA 2% in PBS (MW 150W, 1’). Slides were finally mounted in a 1:1 solution of AF1 antifadent (Citifluor) with PBS, containing calcofluor as a cell wall counterstaining, and imaged by confocal laser scanning microscopy (Zeiss LSM700).

### Cryosections and staining

Freshly sampled stems were kept in −80℃ until sectioning. Leica CM3050S cryotome was set to - 26℃ and 10µm stem cross-sections were obtained, and immediately mounted in a drop of water. Safranin (0.5% in 50% ethanol) and alcian blue (1% in water) were used for staining.

### Atomic force microscopy (AFM)

AFM on stem cross sections was performed and results calculated as previously described (47), with minor set up changes. Stem cross sections from the internode 12 in both WT and *eki* trees were manually dissected, mounted on slides and kept in a humid chamber until analysis. The handmade cuts of approximately 300µm were made using fresh Wilkinson sword razor blades. All slides contained at least 3 WT and 3 *eki* cross-sections, and 3 independently grown clones of WT (14-038-17) or *eki* (14-038-01) trees were used. Trees were grown in Helsinki and transported to Cambridge for analysis. The same experiment was repeated once. AFM JPK Nanowizard was used. The scanning of wood forming tissue was initiated in the inner border of phloem fibers and moved 5 times towards the mature xylem. All sections were scanned left to right and stereomicroscope pictures of the initial and final cantilever position were acquired to later identify the corresponding tissues. Data analysis was performed using JPK Data Processing software (ver. Spm - 4.0.23, JPK Instruments AG, Germany) and Matlab was used to generate the heat maps of Young modulus as well as to do the statistical analysis. The AFM data were collected following the same protocol as previously described (47). Sliced cross-sections were immobilized on glass slides and surrounded by stiff agarose. Measurement of cell wall properties alone was ensured by suppression of turgor pressure by immersion of all cuts in a hypertonic solution a minimum of 20 minutes before measurement (0.55 M mannitol). The following cantilevers were used: “Nano World” (NanoWorld AG Headquarters, Neuchâtel, Switzerland) TL-NCH-20 tips with a spring constant of 10 – 130 N/m (those used were estimated to be 1.5 N/m) with Sphere Tips of a 25 µm radius. All force spectroscopy experiments were performed as previously described; briefly, rigidity of samples was determined as follows: an AFM cantilever loaded with a spherical tip was used to indent the sample over a 100 × 100 µm square area, within the area 100 × 100 µm measurements were made resulting in 10000 force-indentation experiments, each force-indentation experiment was treated with a Hertzian indentation model to extrapolate the apparent Young’s modulus, each pixel in a rigidity map represents Young’s modulus from one force-indentation point.

The maximum force applied by the cantilever was set as to perform a deformation approximately of 2 µm: Set point: 381.323 nN Re.Setpoint: 4.500nN, z-length: 5, z-movement: constant speed, Extend Speed: 200, Sample Rate 8000. In all cases were sample movement or outreach sample height was detected, results were not considered.

### Tensile strength testing of full stems

Three point bending tests were performed to analyze the mechanical properties of the full stems using Texture analyser TA.XT (Stable Micro Systems Ltd., Godalming, Surrey, U.K.). A 5 kg load cell was used. Previously, samples were prepared by selecting internodes of approximately same diameter (2.83∓0,27mm), from the middle height of the stem in WT and *eki*. Samples were dried overnight at 65°C. The measurement procedure was based on descriptions of similar tests by Horvath et al. (2010) and Sindhu et al. (2007), with modifications. As the TA.XTs three point bending rig and probe (HDP/3PB) were too large to be used with these samples, a custom rig and probe with smaller dimensions were constructed. After attaching the fixture and the probe, a calibration was performed. The stem sample was placed on the fixture with a 25mm gap width. The aluminum probe (width 1mm) pushed down the sample from the middle. The speed of the probe was set to 1 mm/s. Three biological replicates and three technical replicates were measured. The sample diameter was 2,83±0,27mm. The Texture Exponent 32 version 6 software recorded the force (g) and distance (mm) data. Hardness is determined as the force (g) required to break the sample. Brittleness is defined as the distance to break (mm). It does therefore indicated how far a sample can be deformed before fracture. Toughness is determined as the slope of these two (mm/g).

### Ploidy estimation by flow cytometry

Samples of ∼1 cm^2^ leaf from in-vitro grown plant material was collected in a pre-chilled clear flat glass petri-dish. The samples were chopped with razor blades ≤60 seconds with slight modification of woody plant buffer(48) (200mM Tris, 4 mM MgCl_2_, 2 mM Na_2_EDTA, 86 mM NaCl, 10 mM sodium metabisulfite, 1% PVP-40, 1% (v/v) Triton X-100. Adjust pH 7.5. Store the buffer at 4°C.) The samples filtered through 40µm nylon mesh filter (Corning^TM^ C431750) and stained with Propidium Iodide (PI) 50µg ml^−1^ simultaneously with RNase at 50µg ml^−1^. Incubate for 1 hour dark on ice. Flow cytometry was done using BD LSR II flow cytometer (BD FACSDiva software 8.0.1) as per instrument user instructions. Long day, greenhouse grown rice leaves samples used as reference genome. The samples were kept on ice throughout the procedure.

### DNA Sequencing

Libraries for genome sequencing were done as described in Salojärvi et al., 2017 (page 26 section 1.3.1.2). The libraries had a insert size of approx. 350 bp. and were paired-end (150 bp+150 bp) sequenced on a NextSeq 500 instrument (Illumina) at the DNA Sequencing and Genomics Laboratory, Institute of Biotechnology, University of Helsinki.

### Genome analysis, linkage mapping and QTL analysis

Mapping data to the genome

The backcross data was analyzed as follows.

First, the individual fastq files were mapped to *B. pendula* genome (20) using bwa mem (49) and sorted bam files were produced using SAMtools (50).

#### Verifying tetraploidy

Then, SAMtools mpileup and a custom script was used analyse allele ratios of variant bi-allelic sites on the parental individuals. Only sites with total sequencing coverage between 12-100 and a minimum allele coverage of 3 were considered as variants. Density estimation and histogram was applied to the allele ratios in R (51).

#### Linkage mapping

Lep-MAP3=LM3 (22) was used to construct linkage map for the backcross data. The LM3 variant calling pipeline was modified to call tetraploid genotypes. In more detail, pileup2posterior.awk script was replaced with pileup2posterior_polyploid.awk and ParentCall2 with ParentCall2Ploidy. The pipeline only considers markers where parents are simplex markers, i.e. containing one different allele like AAAB for tetraploid with some nucleotides A and B.

After the modified variant calling pipeline, the data was analysed as any diploid data with LM3. Markers informative only in mother or father were first clustered into linkage groups using SeparateChromosomes2 with LOD score of 12 (lodLimit) and distortionLod=1 and more markers were added to these groups with JoinSingles2All with LOD 10.

This yielded 28 linkage groups for paternal maps (two per each chromosome of B. pendula) and 31 for maternal (two for chrs 2-14 and 5 for chr1). The 5 groups in chr1 were further grouped into two groups by extending (at most 5kb) the markers in the two paternal linkage groups for chr1 and then running SeparateChromosomes2 on these markers without distortionLod (effectively smaller LOD) parameter. The total number of paternal and maternal markers assigned to these groups were about 1.3M and 1.2M.

The markers in each of these linkage groups were ordered with OrderMarkers2 and Marey maps were plotted in R (Fig S10).

#### QTL analysis

LM3 outputs the segregation pattern for each markers, a custom script was used to calculate the LOD score between this pattern and *eki*/WT pattern of each offspring. Closest pattern matching the phenotype in maternal maps had 12 (out of 60) differences and 16 in paternal. The significance of these associations was calculated in R from binomial distribution with success parameter of 0.5 and using number of unique markers (2740) as the Bonferroni (multiple test) correction term (R: 2740*2*pbinom(12, 60, 0.5)). The most significant region was located at the end of chr9.

#### Genome anchoring

As each *B. pendula* chromosome corresponded to four linkage groups of 56, the *B. pendula* contigs were put to 14 chromosomes based on the map (similarly as in Salojärvi et al., 2017) and placed and ordered based on marker positions. The Marey map (Fig S10) for the resulting genome was constructed using custom R script.

#### Pooled data analysis

To verify the QTL results, the back-cross offspring were analysed as two pooled samples, one pool for all individuals showing WT phenotype and another for *eki*. For both pools and each genomic position the number of nucleotides A, C, G and T were calculated omitting positions where total counts were >1000. Likelihood ratio of the counts coming from one multinomial vs. two (corresponding the pools) multinomial distributions was calculated. Based on a random assignment of phenotypes and calculating the LOD, a LOD limit 13 was used to find significant regions. The most significant region from this analysis was overlapping the QTL region in chr11.

### RNA-Seq analysis

#### Plant materials

Samples of internode 2 (IN2), bark (IN5bark) and xylem (IN5xylem) from internode 5 were collected from four wildtype and mutant named “*Elimäki*” individuals growing in the greenhouse, respectively. All materials were immediately frozen in liquid nitrogen and stored at −80°C for RNA extraction.

#### RNA extraction and sequencing

RNA was extracted as described previously (52); and RNA quality was verified by RNA Nano Chip from Agilent Technologies (US). The 78bp/74bp paired-end sequences were generated using HiSeq2000 platform at the Sequencing and Genomics Laboratory, Institute of Biotechnology, University of Helsinki.

#### Bioinformatics analysis

The raw sequencing data were firstly assessed using FastQC (v0.11.8, http://www.bioinformatics.babraham.ac.uk/projects/fastqc/). The residual ribosomal RNA (rRNA) contamination was filtered using SortMeRNA (v2.1, (53)). The removal of adaptors and low-quality reads and trimming of low-quality nucleotides were performed using Trimmomatic-0.35 (54). The quality of processed reads was then assessed again with FastQC. High quality reads for each sample were mapped to the genome of *B. pendula* (v1.2) with corresponding gene model annotation using HISAT2 2.1.0 (55). Read counts per gene were generated by featureCounts function in the Rsubread package. Transcripts per million (TPM) method was used to normalize reads in each sample, and trimmed mean of M-values (TMM) normalization method was used to equate the overall expression levels of genes between different samples. Differential expression analysis between mutant and wildtype for each type of sample was done by DEseq2 R package (56). DEGs were filtered following the criteria of an absolute value of log_2_fold change > 1 and adjusted *p*-value < 0.05. Gene Ontology (GO) annotations for all genes were obtained as recently described (57), and GO functional enrichment analysis was performed by utilizing clusterProfiler R package (58).

## Supplementary Materials

**Supplementary Data 1:** Candidate genes in EKI locus (Chr11) and RNASeq based expression profiles.

**Supplementary Data 2:** BC_1_ individuals sequencing ID and phenotyping information.

**Supplementary Data 3:** Complete analysis of DEGs from RNASeq.

### Supplementary Note 1: Quantitative trait locus (QTL) mapping combined with RNASeq identifies DEGs at the EKI locus

The identified mapping window in chromosome 11, contains 324 annotated genes in *B. pendula* genome (candidate list). Given that phenotypes linked to recessive mutations frequently result in transcriptional downregulation of the causative gene, we next explored the expression profile of genes this subset of genes. First, we relaxed the cutoff of DEGs by including all genes where the fold change is significantly different in WT vs. *eki* (*p* adj<0.05). From our mapping window we found that a total of 23 genes are differentially regulated in all RNASeq samples (Fig S4b, **Supplementary Data 1**).15/23 genes were downregulated in all *eki* samples but only 4 genes were had barely detectable levels of gene expression in *eki* (< 1 Avg. TPM), and higher expression in all WT samples. These four genes are Bpev01.c2969.g0002, Bpev01.c0954.g0001, Bpev01.c1790.g0003, and Bpev01.c0306.g0009. Based on sequence homology, Bpev01.c2969.g0002 encodes for a member of the hAT transposon superfamily. Both, Bpev01.c0954.g0001 and Bpev01.c1790.g0003 are orthologues of the Arabidopsis PR5-LIKE RECEPTOR KINASE (AT5G38280) which has been associated to plant defense (59). Finally, Bpev01.c0306.g0009 shares partial similarity with *Quercus suber* putative phospholipid-transporting ATPase 9 (LOC111987270). While these candidates are genetically linked the *eki* phenotype and could potentially explain it, further analysis and experiments are required to connect them with the *eki* phenotype.

**Table S1:** The effect of stem weight manipulations represented by the proportional changes in stem diameters. Values in the table represent the proportional change in final vs. initial (T4/T1) diameter measurements along the stem. Five trees were used in each of the four different scenarios. WT and *eki* were intercalated within each experiment. The diameters were measured along the stem: in the base (last four internodes, D1-D4), 50% height internode (D5), 62.5% height internode (D6) and 75% height internode (D7).

| Treatments | WT (diameter T4/diameter T1) |  |  |  |  |  |  | *eki* (diameter T4/diameter T1) |  |  |  |  |  |  |
| --- | --- | --- | --- | --- | --- | --- | --- | --- | --- | --- | --- | --- | --- | --- |
|  | D1 | D2 | D3 | D4 | D5 | D6 | D7 | D1 | D2 | D3 | D4 | D5 | D6 | D7 |
| No weight tree #1 | 1,539 | 1,579 | 1,598 | 1,482 | 1,496 | 1,699 | 1,764 | 1,859 | 1,695 | 1,564 | 1,521 | 1,455 | 1,422 | 1,529 |
| No weight tree #2 | 1,528 | 1,481 | 1,459 | 1,443 | 1,602 | 1,529 | 1,760 | 1,494 | 1,491 | 1,413 | 1,466 | 1,406 | 1,500 | 1,545 |
| No weight tree #3 | 1,539 | 1,438 | 1,402 | 1,435 | 1,428 | 1,516 | 1,536 | 1,433 | 1,468 | 1,455 | 1,443 | 1,642 | 1,521 | 1,698 |
| No weight tree #4 | 1,450 | 1,407 | 1,420 | 1,397 | 1,379 | 1,500 | 1,508 | 1,588 | 1,540 | 1,547 | 1,604 | 1,541 | 1,496 | 2,484 |
| No weight tree #5 | 1,533 | 1,479 | 1,479 | 1,455 | 1,615 | 1,636 | 1,811 | 1,635 | 1,589 | 1,548 | 1,556 | 1,538 | 1,717 | 1,681 |
| Weight Supp tree #1 | 1,516 | 1,369 | 1,424 | 1,339 | 1,537 | 1,437 | 1,770 | 1,495 | 1,573 | 1,502 | 1,571 | 1,527 | 1,597 | 1,596 |
| Weight Supp tree #2 | 1,425 | 1,369 | 1,394 | 1,382 | 1,432 | 1,438 | 1,766 | 1,508 | 1,442 | 1,357 | 1,512 | 1,477 | 1,640 | 1,694 |
| Weight Supp tree #3 | 1,505 | 1,536 | 1,480 | 1,474 | 1,640 | 1,873 | 2,340 | 1,737 | 1,554 | 1,574 | 1,611 | 1,483 | 1,558 | 1,932 |
| Weight Supp tree #4 | 1,323 | 1,311 | 1,294 | 1,314 | 1,592 | 1,638 | 1,584 | 1,343 | 1,512 | 1,401 | 1,532 | 1,622 | 1,803 | 1,780 |
| Weight Supp tree #5 | 1,425 | 1,473 | 1,536 | 1,402 | 1,454 | 1,521 | 1,760 | 1,618 | 1,672 | 1,464 | 1,421 | 1,802 | 2,097 | 2,038 |
| Weight Free tree #1 | 1,466 | 1,371 | 1,489 | 1,317 | 1,505 | 1,672 | 1,768 | 1,656 | 1,492 | 1,513 | 1,506 | 1,452 | 1,605 | 1,672 |
| Weight Free tree #2 | 1,522 | 1,520 | 1,485 | 1,523 | 1,628 | 1,717 | 1,961 | 1,543 | 1,531 | 1,447 | 1,621 | 1,109 | 1,648 | 1,915 |
| Weight Free tree #3 | 1,463 | 1,588 | 1,584 | 1,445 | 1,621 | 1,961 | 1,937 | 1,066 | 1,446 | 1,613 | 1,663 | 1,686 | 1,692 | 1,886 |
| Weight Free tree #4 | 1,553 | 1,460 | 1,487 | 1,429 | 1,475 | 1,476 | 1,625 | 1,437 | 1,393 | 1,460 | 1,680 | 1,670 | 1,838 | 1,874 |
| Weight Free tree #5 | 1,745 | 1,764 | 1,833 | 1,556 | 1,673 | 1,734 | 1,894 | 1,551 | 1,498 | 1,484 | 1,476 | 1,710 | 1,624 | 1,859 |
| Weight Pulling tree #1 | 1,643 | 1,472 | 1,464 | 1,451 | 1,636 | 1,662 | 1,753 | 1,533 | 1,626 | 1,583 | 1,549 | 1,623 | 1,684 | 1,730 |
| Weight Pulling tree #2 | 1,456 | 1,428 | 1,419 | 1,514 | 1,688 | 1,743 | 1,739 | 1,662 | 1,599 | 1,524 | 1,552 | 1,650 | 1,572 | 1,734 |
| Weight Pulling tree #3 | 1,479 | 1,445 | 1,456 | 1,458 | 1,565 | 1,602 | 1,589 | 1,418 | 1,429 | 1,441 | 1,543 | 1,466 | 1,519 | 1,757 |
| Weight Pulling tree #4 | 1,414 | 1,403 | 1,362 | 1,392 | 1,553 | 1,684 | 1,871 | 1,648 | 1,601 | 1,527 | 1,688 | 1,575 | 1,671 | 1,812 |
| Weight Pulling tree #5 | 1,422 | 1,387 | 1,133 | 1,499 | 1,509 | 1,737 | 1,708 | 1,598 | 1,487 | 1,617 | 1,520 | 1,538 | 1,319 | 1,708 |

Stem positions

**Table S2:** Segregation Test. The null hypothesis (H_0_) is an expected segregation of 1:1. The observed phenotypes frequency indicates that there is no evidence to reject H_0_, (Chi square *p* value is ≥ 0.05).

| Class | Observed | Expected |
| --- | --- | --- |
| *eki* | 27 | 30 |
| WT | 33 | 30 |
| Total | 60 | 60 |
|  | CHISQ.test pvalue | 0,438578026 |

**Figure S1.**
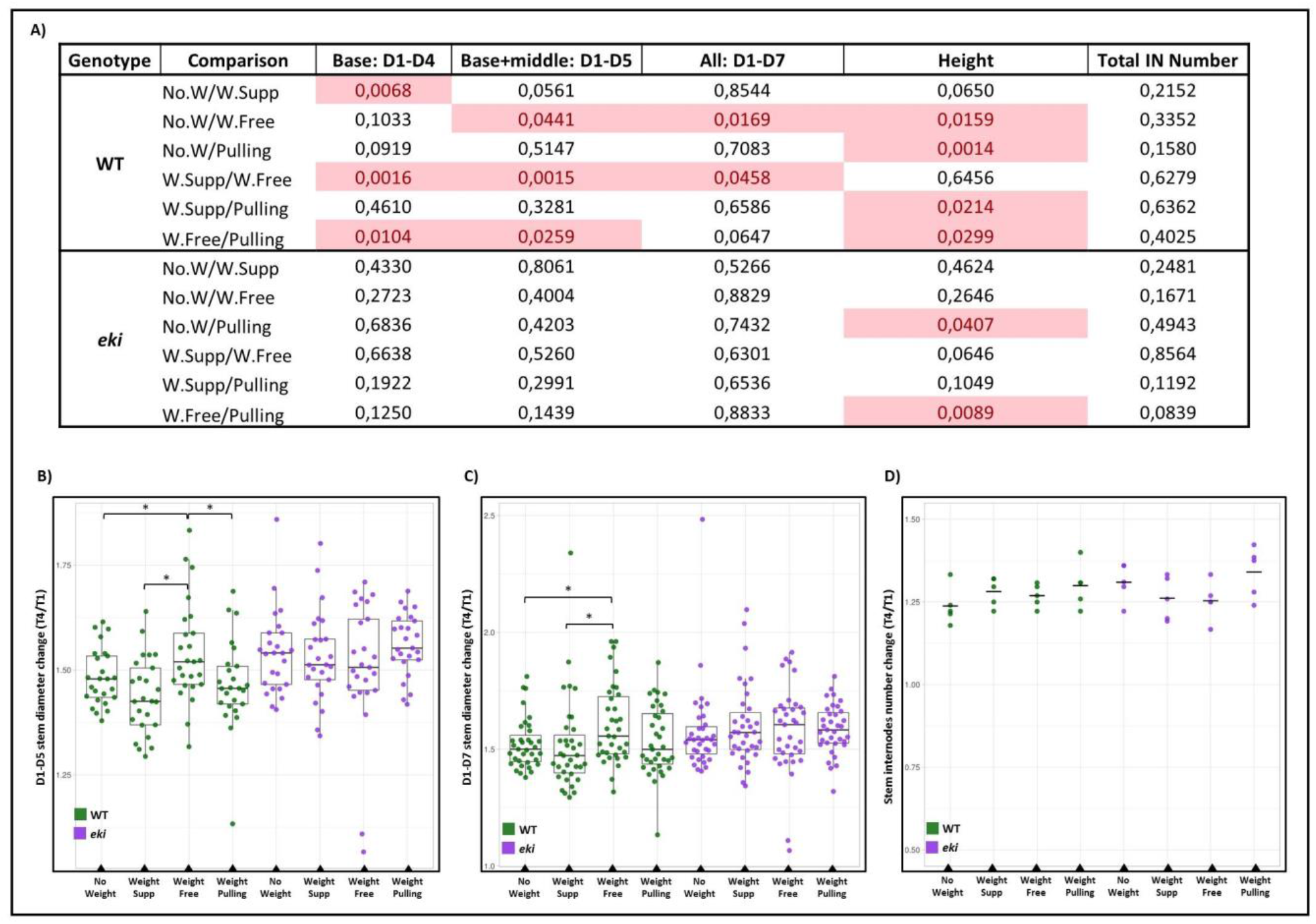
Vertical mechanical manipulations in WT vs. *eki*. **A)**Table of *p* values obtained from two-tailed Student t-test comparing: Scenario 1 (No.W = no weight added), Scenario 2 (W.Supp = weight added on supported stems), Scenario 3 (W.Free = weight added on semi-free stems), Scenario 4 (Pulling = stems pulled up from the top). **B)** Stem diameter change in final vs. initial timepoints (T4/T1) on the combined internodes (D1-D5). **C)** Stem diameter change in final vs. initial timepoints (T4/T1) on the combined internodes (D1-D7). **D)** Internode numbers change in final vs. initial timepoints (T4/T1). Each experimental setup consisted in 5 biological replicates per treatment. Asterisk indicate statistically significant differences *p* ≤0.05 after Student t-test.

**Figure S2:**
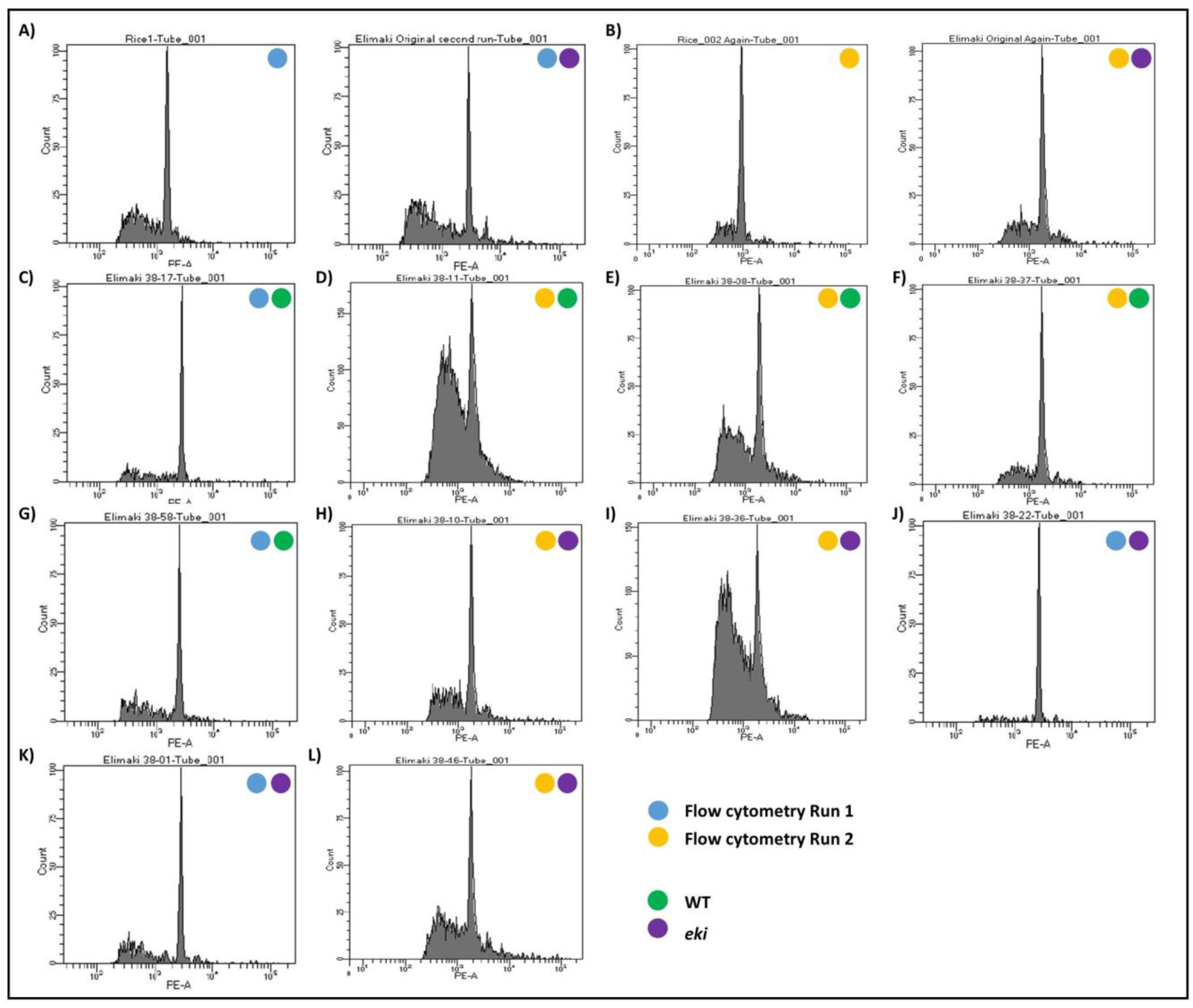
Flow cytometry ploidy analysis. The analysis was performed in two batches, each with the controls: rise and Elimäki Original (EO) **A) Left:** *Oryza sativa* was used as a diploid genome control (430Mb). **Right:** EO shows a peak that doubles the genome size of rice, similar results were reported before (20). **B)** Second run of the same controls shown in (A). Remaining plots correspond to individuals from the BC_1_ population showing peaks of genome sizes similar to their corresponding EO control. **C-G) Wild type trees**: #38-17, #38-11, #38-09, #38-37, #38-58; **H-L) *eki* trees**: #38-10, #38-36, #38-22, #38-01, #38-46.

**Figure S3.**
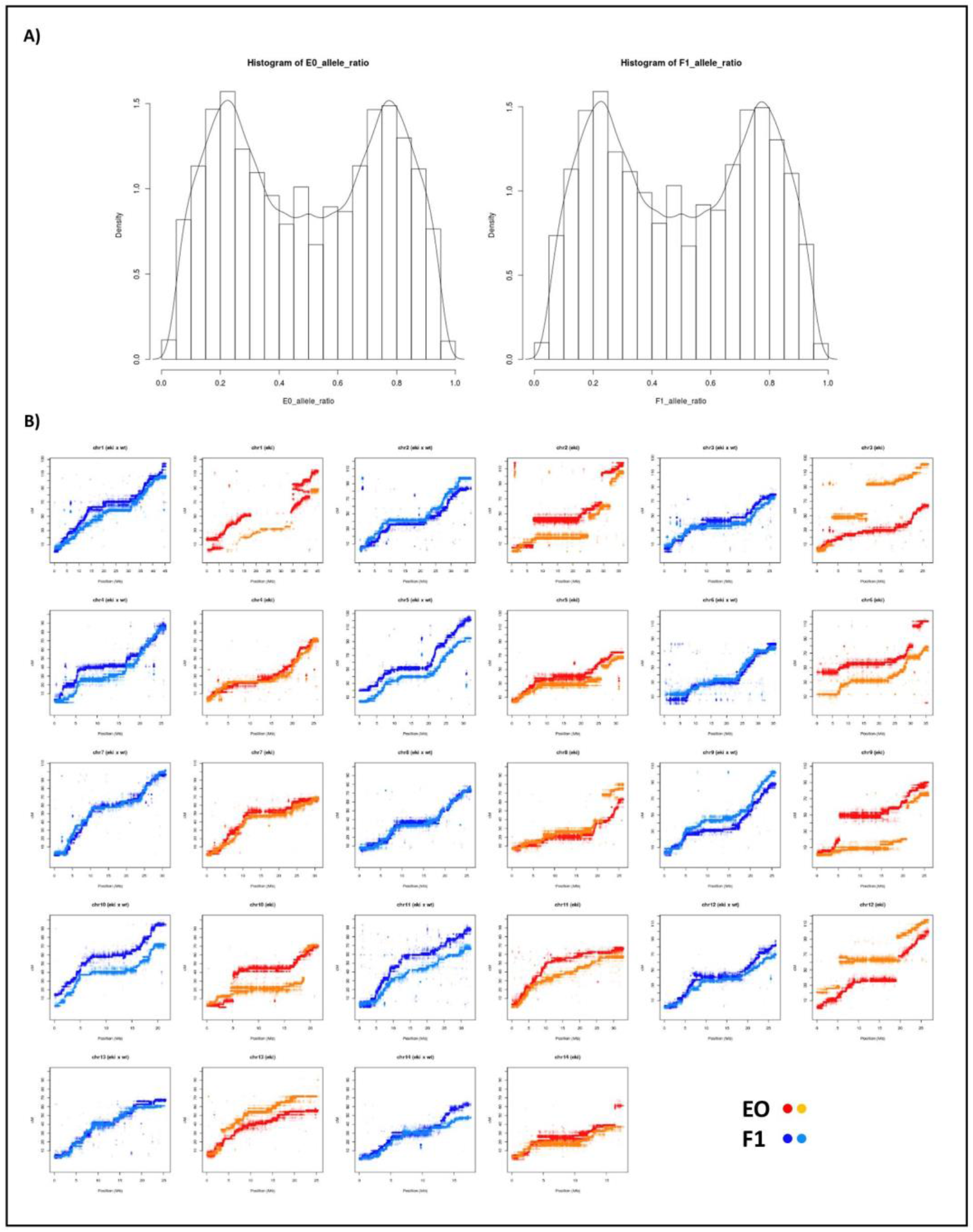
**A)** Allele ratios at bi-allelic variants for EO and F1 individuals. Peaks at about 1/4 and 3/4 correspond to bi-allelic tetraploid markers of types AAAB and ABBB, thus supporting tetraploid fashion of these individuals. **B**) Marey maps for all 56 linkage groups. Note that each *B. pendula* chromosome has 4 linkage groups, two maternal from EO (red and orange) and two paternal from F1 (light and dark blue).

**Figure S4.**
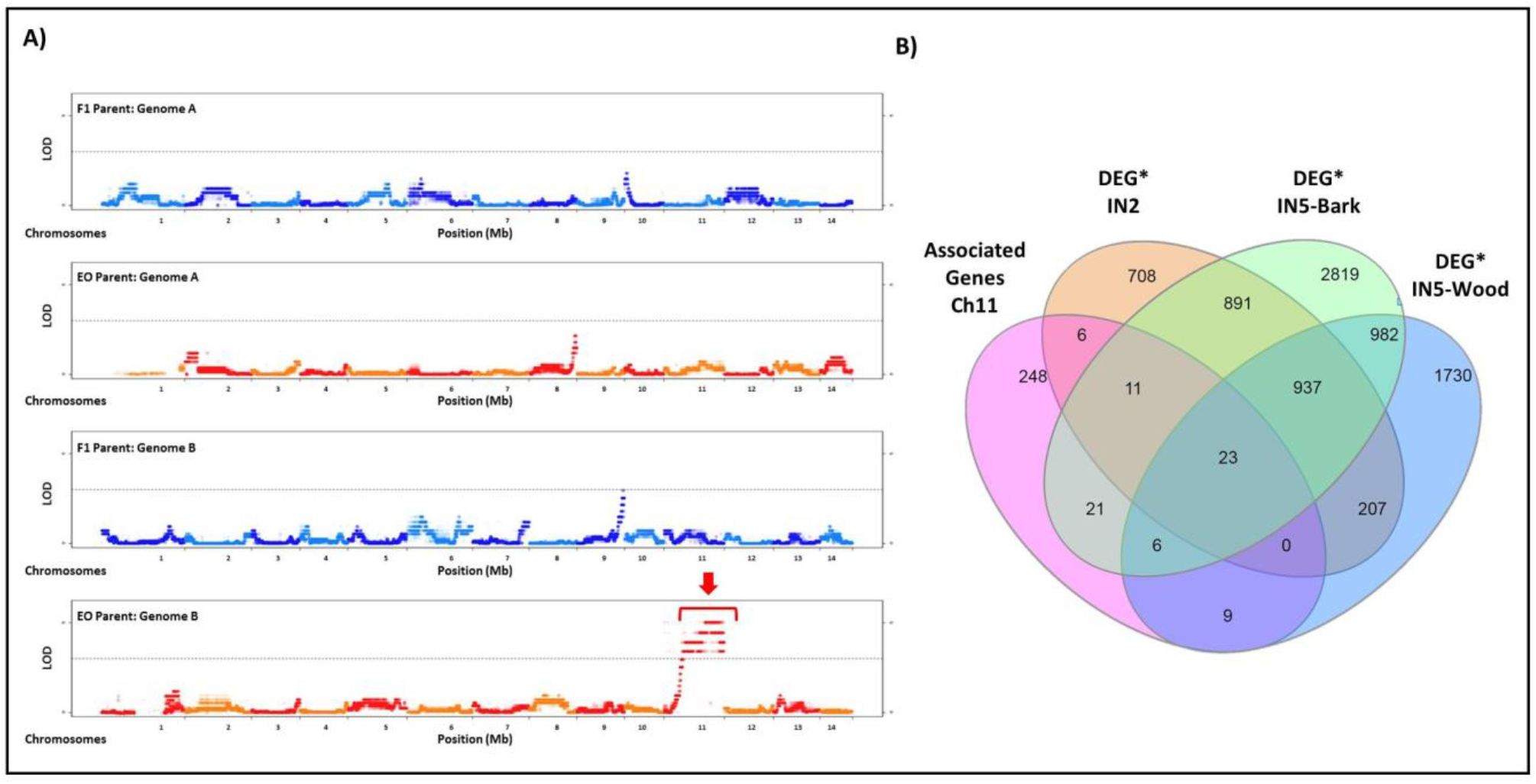
Identification of candidate genes by QTL mapping and RNASeq. **A)** Manhattan Plot showing the association of polymorphic markers to the parents F1 (WT phenotype) and EO (eki phenotype). The tetraploid genome was separated in Genome A and Genome B. The horizontal dotted line indicates the significance cutoff for the identified associations. **B)** Venn diagram with Candidate Genes from mapping window in the chromosome 11 (Chr11) and genes with significant differential expression (*p* adj<0.05) within each sample.

**Figure S5:**
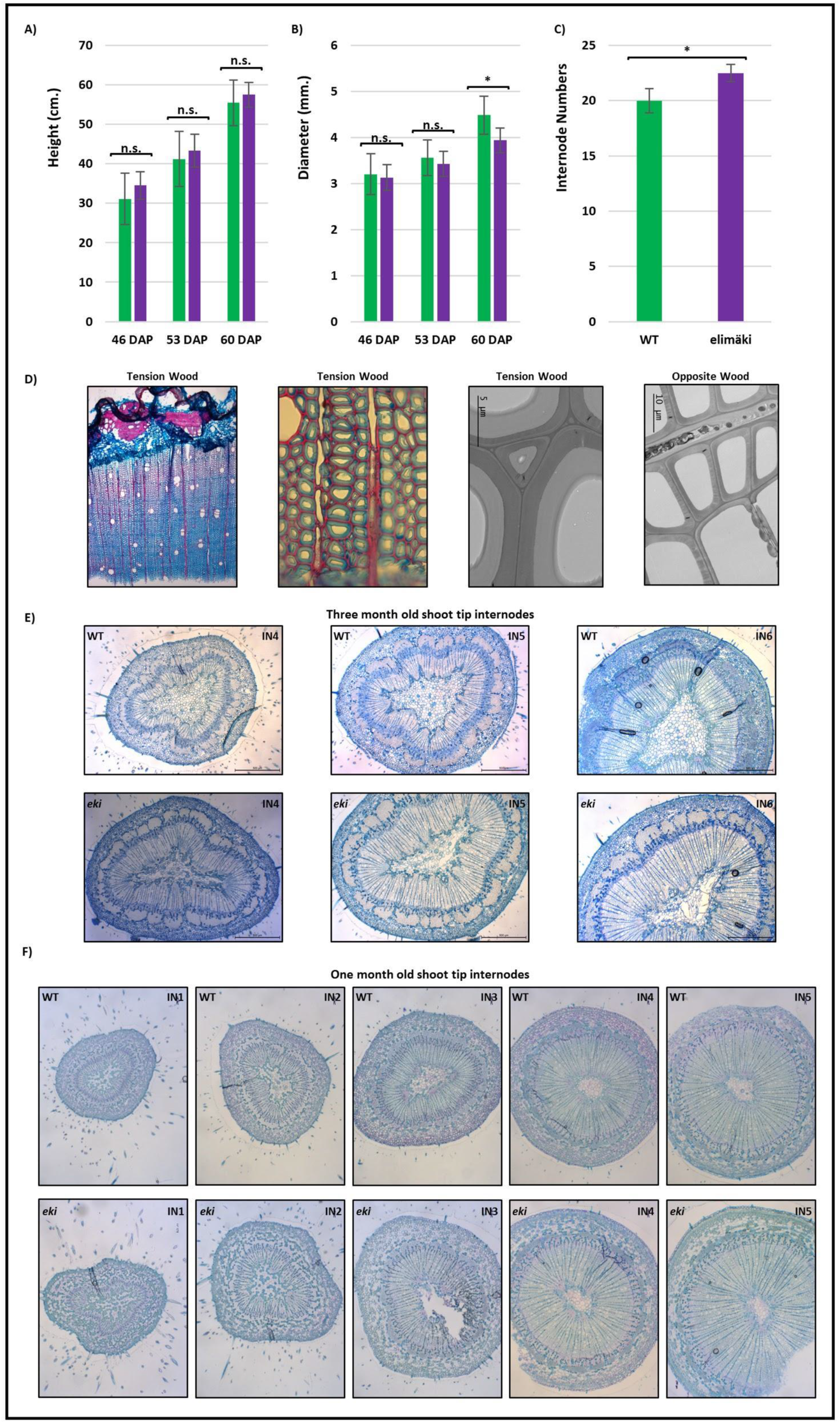
Developmental phenotypes before and after collapse. Before collapsing WT vs. *eki* growth dynamics in 46, 53 and 60 days after potting (DAP) represented by the total height **(A)** and diameter at the fixed position of 10cm above base **(B)**. WT, n = 9; *eki*, n = 6 trees. **C)** Total internode numbers of trees before collapse. Asterisk indicate statistical significant differences (Student-t test) comparing WT vs. *eki*. n.s. = non-significant. **D)** Two months after collapse, the anatomy of main stem in *eki* displays a large accumulation of tension wood as revealed in cryosections stained with alcian blue and safranin. TEM images reveal G-layers in the inner side of secondary cell walls in tension wood samples. **E)** Cross-sectioned stained with toluidine blue show the developmental progression of internodes at the shoot apex in three months old trees, *eki* cross-sections revealed a larger xylem expansion. **F)** Cross-sectioned stained with toluidine blue show the developmental progression of internodes at the shoot apex in one month old trees, *eki* cross-sections revealed a larger xylem expansion (quantification in **Fig. 3**).

**Figure S6:**
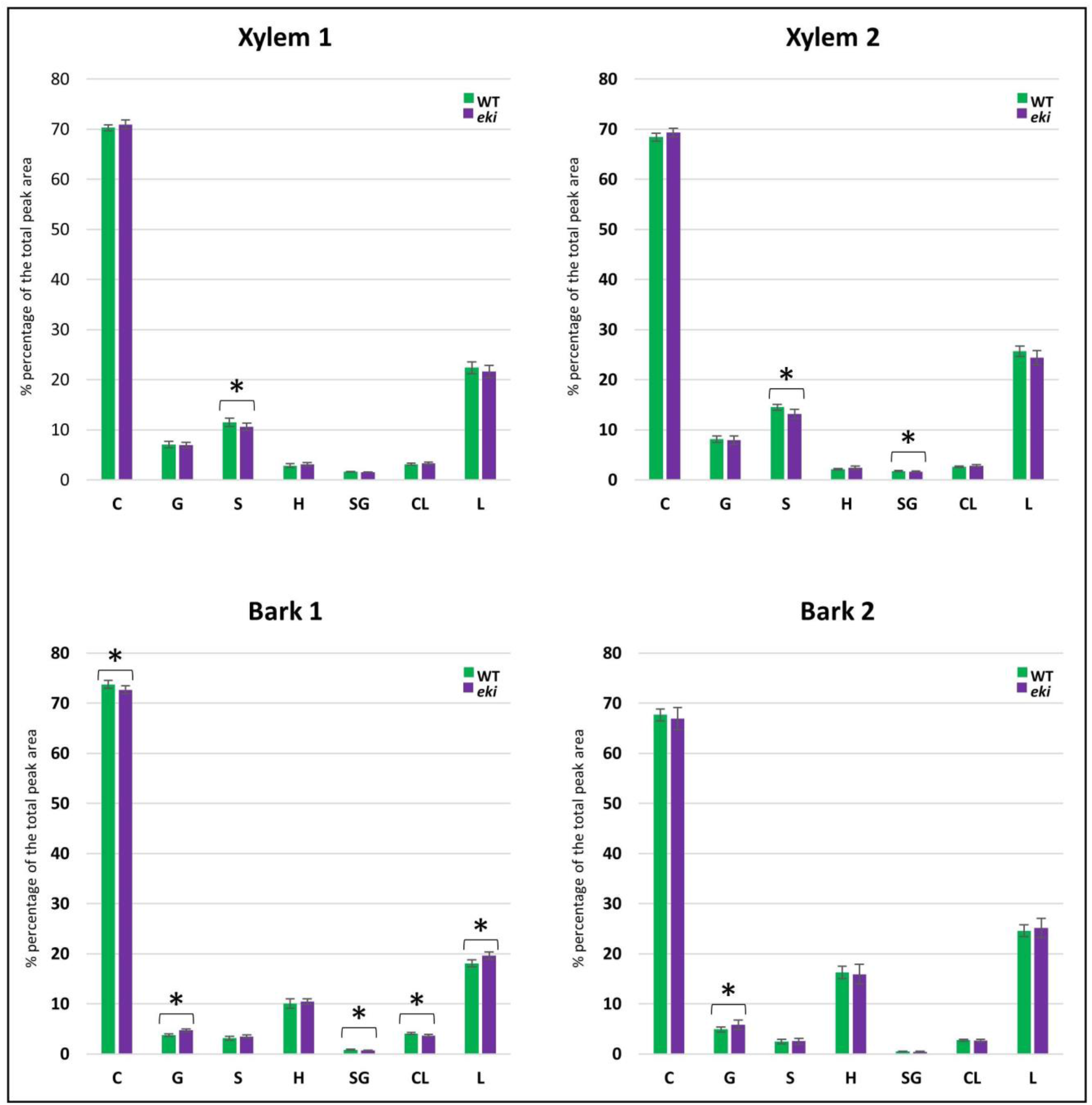
Pyrolysis-GC/MS analysis of xylem and bark from WT and *eki*. Comparison of identified peak areas, values are percentage of the total peak area. **Xylem 1 and Bark 1**: Xylem and bark tissues from internodes 6-7-8. **Xylem 2 and Bark 2:** Xylem and bark tissues from internodes 9-10-11. **C: combined peaks attributable to carbohydrates**, **G: peaks attributable to** guaiacyl lignin, **S: peaks attributable to** syringyl lignin, **H: peaks attributable to** *p-*hydroxyphenyl lignin, **SG:** S/G ratio, **CL:** C/L ratio, **L: combined peaks attributable to** lignin. Bars represent the average value and error bars indicate the standard deviation of 3 biological replicates and 3 technical replicates. (*) Asterisk indicate statistically significant differences (Student t-test) comparing WT vs. *eki*.

**Figure S7:**
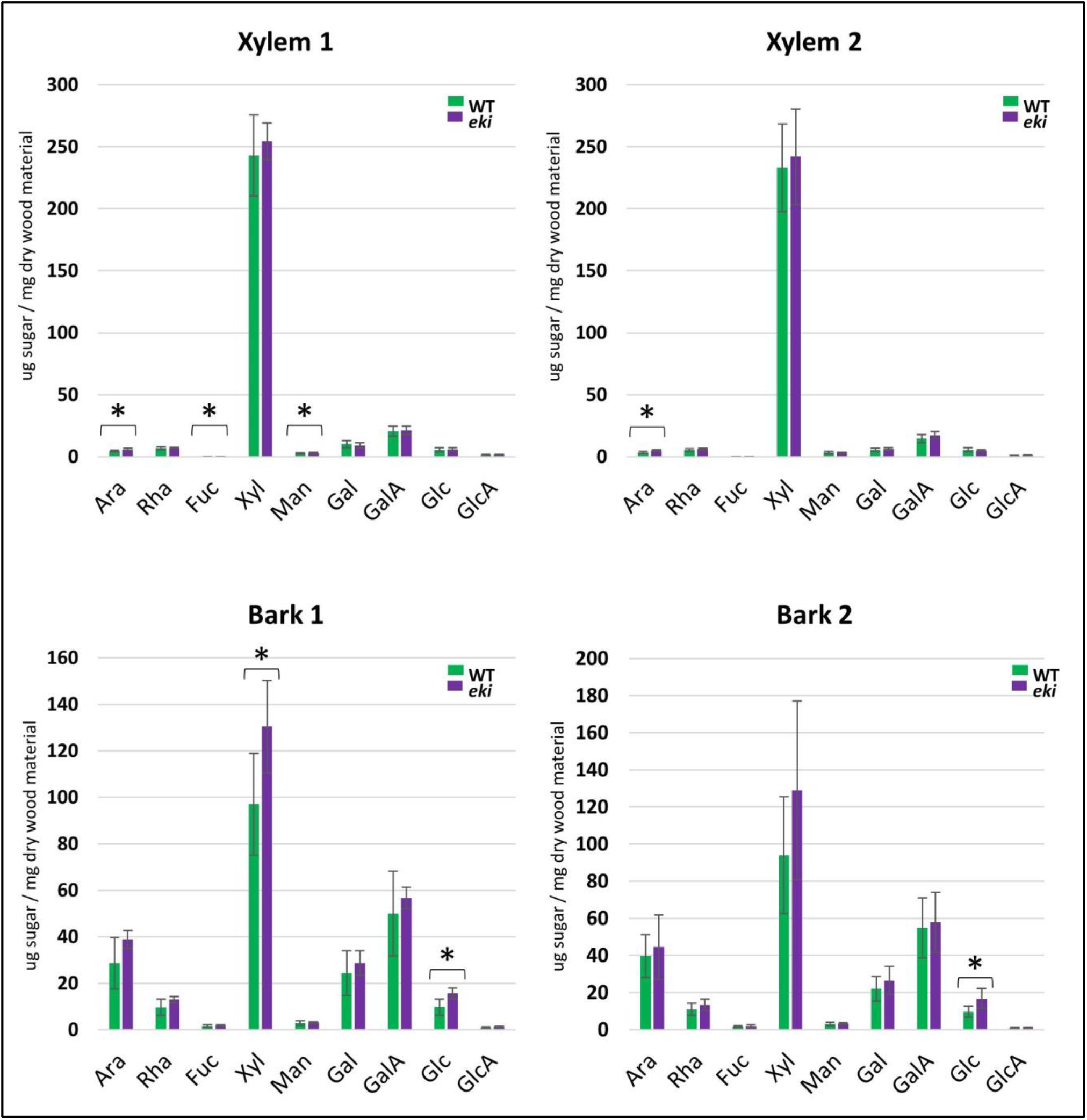
Monosaccharide composition analysis of AIR by TMS derivatization with GC/MS. **Xylem 1 and Bark 1:** Xylem and bark tissues from internodes 6-7-8. **Xylem 2 and Bark 2:** Xylem and bark tissues from internodes 9-10-11. **Ara**: arabinose, **Rha**: rhamnose, **Fuc:** Fucose, **Xyl**: xylose, **Man**: mannose, **Gal**: galactose, **GalA**: galacturonic acid, **Glc**: glucose, **GlcA**: glucuronic acid. Bars represent the average value and error bars indicate the standard deviation of 3 biological replicates and 2 technical replicates. (*) Asterisk indicate statistically significant differences (Student t-test) comparing WT vs. *eki*.

**Figure S8:**
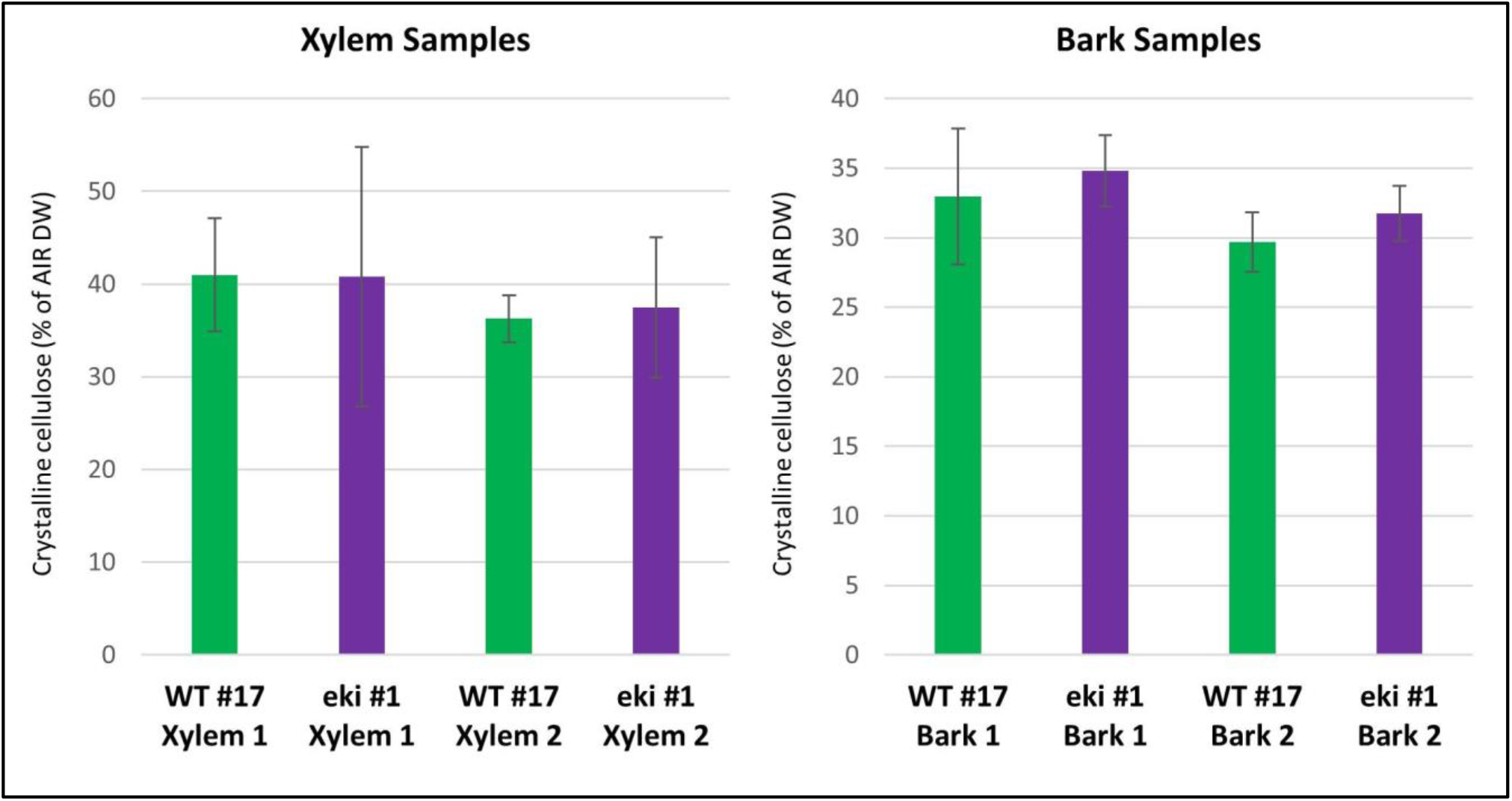
Updegraff cellulose determination of WT and *eki*. **Xylem 1 and Bark 1:** Xylem and bark tissues from internodes 6-7-8. **Xylem 2 and Bark 2:** Xylem and bark tissues from internodes 9-10-11. Bars represent the average value and error bars indicate the standard deviation of 3 biological replicates and 3 technical replicates. No statistically significant differences (*p* <0.05) were found in Student t-test WT vs. *eki* comparisons.

**Figure S9:**
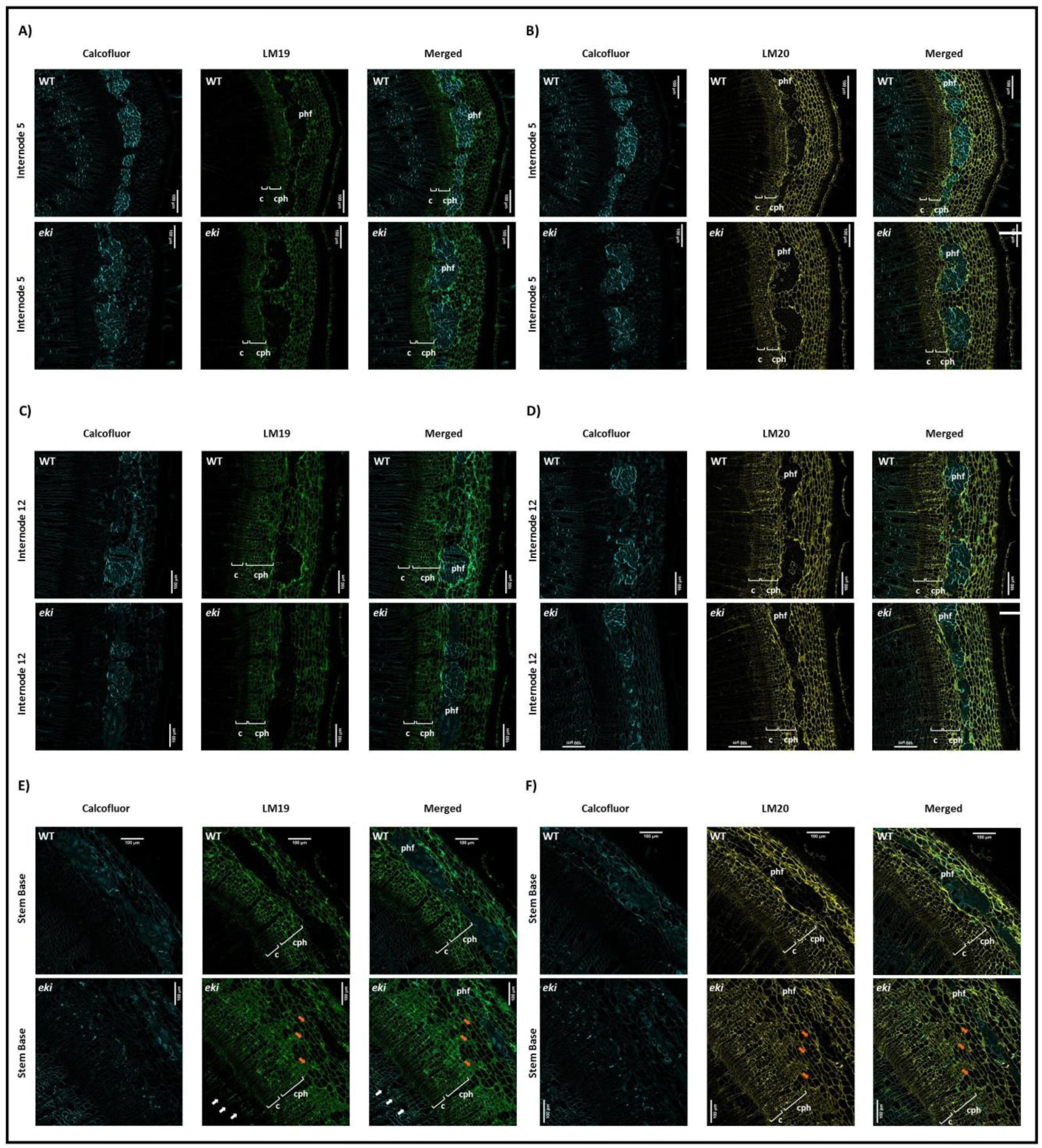
Developing xylem and phloem tissues retain PCW characteristics in *eki*. Immunolocalization of stem cross in internode 5 **(A, B)**, internode 12 **(C, D)** and in the oldest internode at the stem base **(E, F)**. Images correspond to the calcofluor white counterstaining (cyan), and LM 19 (green) or LM 20 (yellow) immunolocalization. Brackets in the figures correspond to: cambium (c), conductive phloem (cph). Phloem fibers (phf). White arrows indicate the LM19 labeling in xylem. In WT trees the conductive phloem connects cambium and phloem fibers, but extra parenchymatous cells labeled by LM19 and LM20 can be found in *eki* (orange arrows).

**Figure S10.**
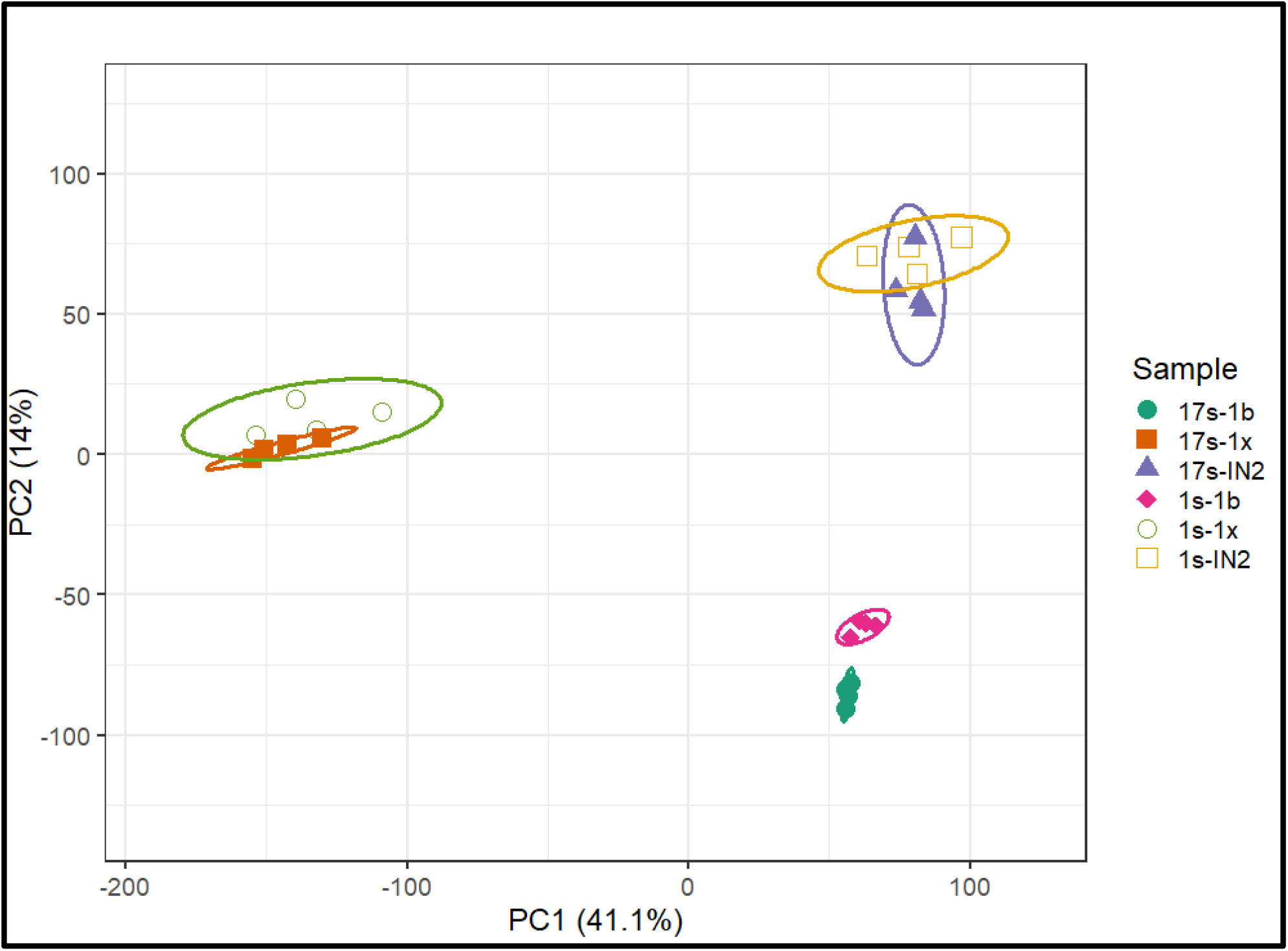
**Principal component analysis of transcriptome profiles for four biological replicates of samples:** 17s = WT, 1s = *eki*, IN2=internode 2, 1b = Internode 5 bark, 1x = Internode 5 wood. Samples group according to their tissue type.

**Figure S11:**
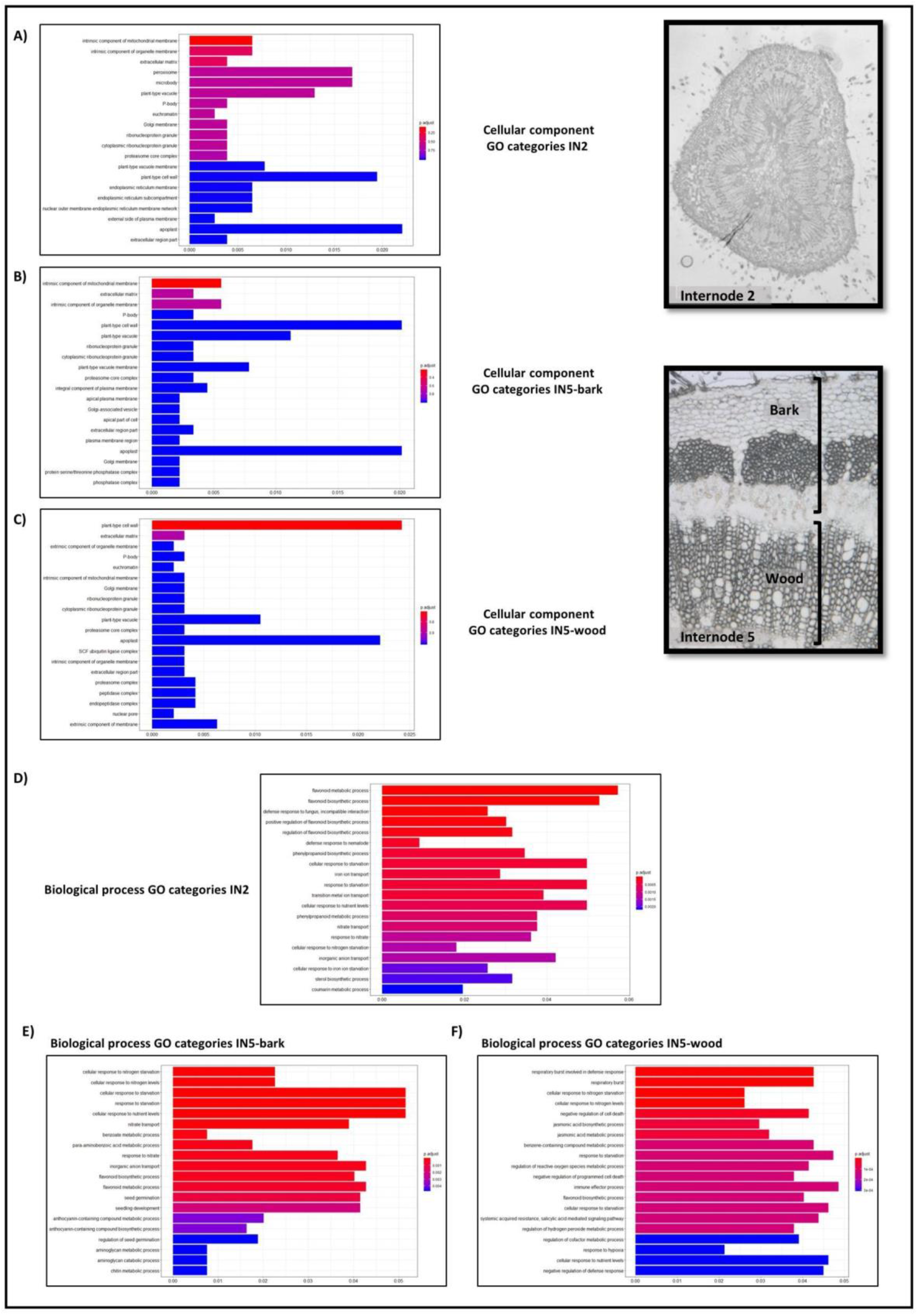
GO enrichment analysis of DEGs in WT vs. *eki*. A-C) GO categories associated to cellular component in A) internode 2, B) IN5-bark and C) IN5- wood. D-F) GO categories associated to biological process in D) internode 2, E) IN5-bark and F) IN5-wood. Microscopic pictures are mere representative figures of the analyzed tissues.

